# Vagus Nerve Stimulation Ameliorates Epilepsy via a Gut Microbiota–Butyrate–Vagus Nerve–Brain Axis

**DOI:** 10.64898/2026.02.20.707138

**Authors:** Long Zeng, Junjie Zou, Yuting Xu, Jingjing Kong, Qingshan Xiao, Zizhang Cheng, Minghan Xie, Yuhui Wan, Anbo Zhang, Yunhao Luo, Yuhan Huang, Yanjuan Ye, Jine Yuan, Shan Xue, Xizhong Zhou, Peihua Cao, Zhaohua Zhu, Tao Xie, Haitao Sun, Yanwu Guo

## Abstract

Vagus nerve stimulation (VNS) is an established therapy for drug-resistant epilepsy, yet its mechanism remains partially understood. Here, through an integrated clinical–translational approach, we demonstrate that VNS modulates the gut microbiome and alter the level of butyrate to reduce seizure susceptibility via vagus nerve–dependent signaling. In a longitudinal cohort of epilepsy patients, VNS enriched short-chain fatty acid–producing bacteria, including *Akkermansia muciniphila*, *Prevotella*, *Alistipes*, and *Odoribacter*. Among clinical responders, fecal butyrate levels decreased concomitantly with an increase in plasma butyrate concentrations. Fecal microbiota transplantation from post-VNS patients into mice reduced seizure severity and improved gut barrier integrity. In a murine model of epilepsy, VNS similarly increased the abundance of Akkermansia and plasma butyrate concentrations, decreased fecal butyrate levels, and abolished its antiseizure effects upon vagotomy. Either colonization with *Akkermansia muciniphila* or exogenous butyrate supplementation was sufficient to reduce seizures and anxiety-like behavior, effects that also required intact vagal signaling. Our findings define a microbiota–butyrate–vagal pathway essential for the therapeutic effects of VNS, highlighting the gut–brain axis as a target for epilepsy treatment.

**GRAPHICAL ABSTRACT:** 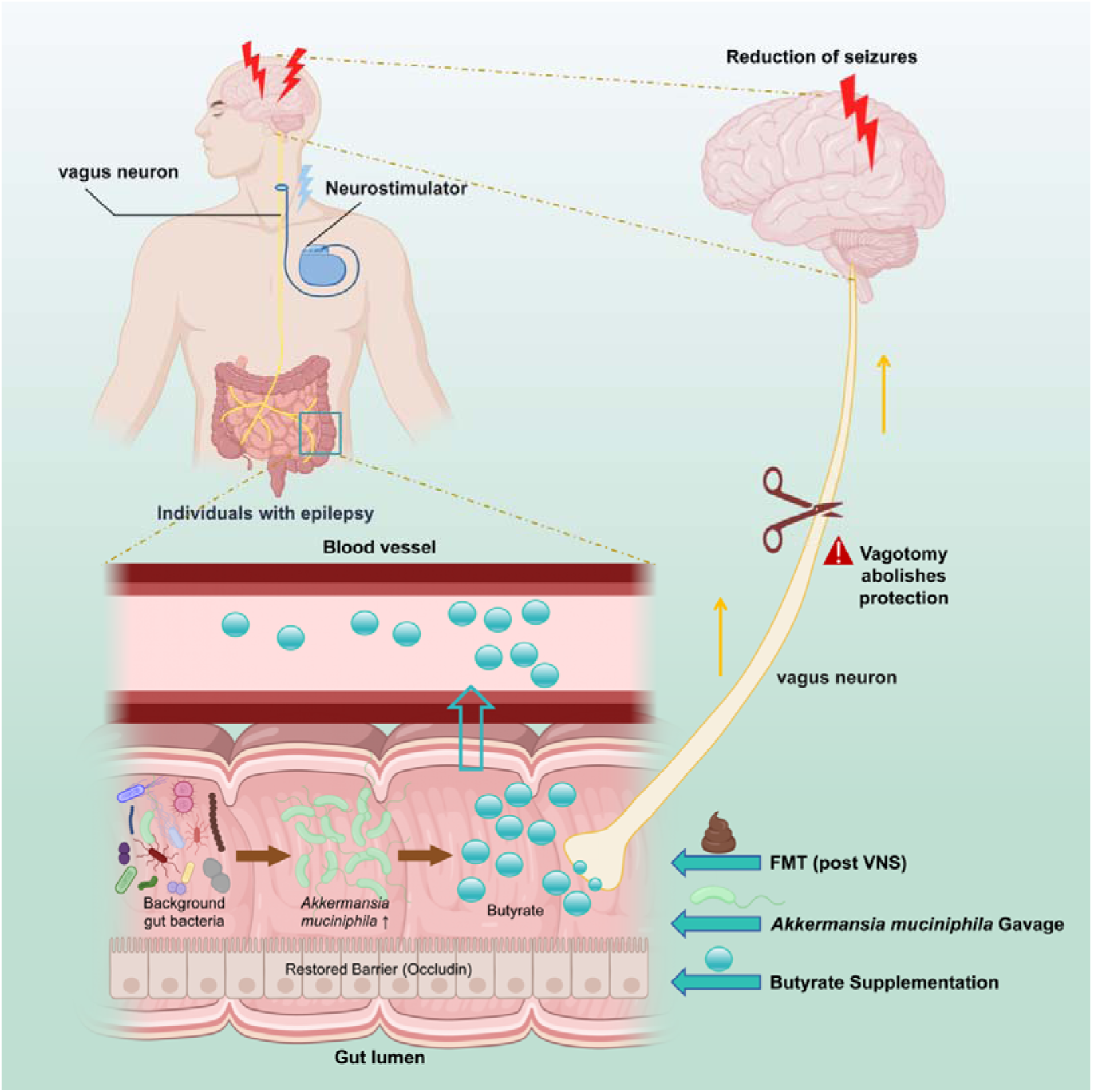

**HIGHLIGHTS:**

- Vagus nerve stimulation (VNS) remodels the gut microbiota and alter the level of butyrate.
- VNS enriches short-chain fatty acids-producing bacteria, specifically *Akkermansia muciniphila*, in both humans and mice.
- Transplantation of VNS-conditioned microbiota confers seizure protection and restores gut barrier integrity.
- *Akkermansia muciniphila* or exogenous butyrate is sufficient to reduce seizures via a vagus nerve-dependent pathway.

## INTRODUCTION

Vagus nerve stimulation (VNS) is an established neuromodulation therapy for drug-resistant epilepsy, proven to effectively reduce seizure frequency in patients(Clifford et al., 2024; Dinger et al., 2025). Despite its clinical efficacy, the precise mechanisms through which VNS exerts its anti-seizure effects remain incompletely understood(Elkattawy et al., 2023; Vitale et al., 2023). Traditional perspectives have largely focused on its direct and indirect neuronal projections modulating excitability within brainstem, thalamic, and cortical networks(Elkattawy et al., 2023; Vitale et al., 2023). In recent years, the emerging concept of the gut–brain axis has revealed a critical role for the gut microbiota in the pathophysiology and treatment of neurological disorders(Al-Beltagi et al., 2023; Li et al., 2025; Z. Liu et al., 2025). Gut microbial communities and their metabolites, particularly short-chain fatty acids (SCFAs) such as butyrate, can influence central nervous system function via immune, endocrine, and neural pathways, and have been closely linked to seizure susceptibility and severity(Paw Owska et al., 2025; You et al., 2025).

Notably, the vagus nerve serves as a major bidirectional communication pathway within the gut–brain axis(Anaclerio et al., 2024; Fernández-Calvet et al., 2024; Liu et al., 2022). Clinical and preclinical evidence suggests that VNS can modulate intestinal motility, permeability, and local immune milieu(Huang et al., 2023; Kawasaki et al., 2011; Vitale et al., 2024). However, whether VNS ameliorates epilepsy by remodelling the gut microbial ecosystem remains systematically unexplored. Furthermore, specific beneficial microbes, such as *Akkermansia muciniphila* (AKK), have been associated with metabolic health, gut barrier integrity, and neuroprotection(Ma et al., 2025; Shi & Li, 2025), yet their potential involvement in VNS-mediated therapeutic effects and their dependence on vagal signaling is still unknown.

Building on this background, this study aimed to systematically investigate whether VNS alleviates epilepsy via a vagus nerve–mediated gut–brain pathway. We first longitudinally characterized changes in gut microbiota composition and metabolomic profiles (with emphasis on SCFAs) in a cohort of patients with drug-resistant epilepsy before and after VNS treatment, correlating these changes with clinical outcomes. Subsequently, through a series of animal experiments-including fecal microbiota transplantation (FMT), VNS intervention, specific probiotic (AKK) colonization, exogenous butyrate supplementation, and vagotomy-we causally validated the essential role of gut microbiota and their metabolites in the anti-seizure effects of VNS, and established the indispensability of vagus nerve signaling in this pathway. Our work provides a novel microbiota–metabolite perspective on the mechanism of VNS and offers a theoretical foundation for microbiota-targeted interventions in epilepsy.

## RESULTS

### VNS efficacy correlates with gut AKK enrichment and butyrate level changes in epilepsy patients

We enrolled 11 patients with drug-resistant epilepsy scheduled for VNS implantation to examine how VNS alters the gut microbiota and metabolome. The cohort included 6 males (54.5%) and 9 children (81.8%), with a median age of 9.2 [IQR 4.2–14.5] years. Fecal and plasma samples were collected before VNS and at 6 and 12 months post-implantation, with no changes in antiseizure medications during follow-up. Seizure frequency (baseline set as 100%) decreased significantly at 6 and 12 months post-VNS (Figure 1B). Six patients achieved ≥50% seizure reduction and were classified as responders. Seizure severity, assessed by the National Hospital Seizure Severity Scale (NHS3), also declined significantly at both time points (Figure 1B). Individual seizure frequencies and NHS3 scores are detailed in Supplementary Table 1.

**Figure 1.**
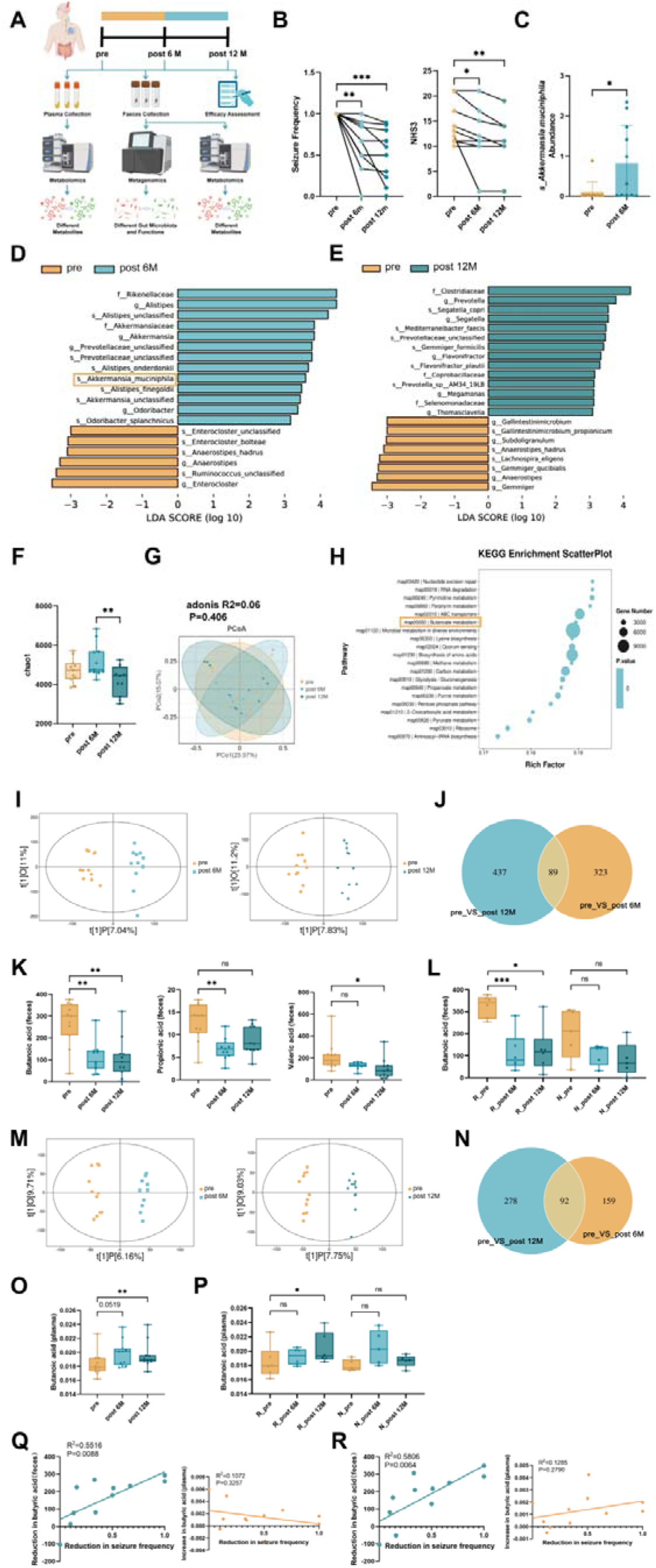
VNS Remodels the Gut Microbiota and Modulates Butyrate level in Patients with Drug-Resistant Epilepsy. **(A)** Study design and flowchart. Plasma and fecal samples were collected from epilepsy patients (n=11) at baseline (Pre-VNS), 6 months, and 12 months post-VNS implantation for metagenomic and metabolomic analyses. **(B)** Longitudinal clinical outcomes showing significant reductions in seizure frequency (left) and severity (NHS3 score, right) at 6 and 12 months post-VNS. **(C, D, E)** LEfSe analysis of metagenomic sequencing data identifying differentially abundant taxa. VNS treatment significantly enriched *Akkermansia muciniphila* at post-operative time points. **(F, G)** Alpha diversity (Chao1 index, **G**) and beta diversity (PCoA based on Bray-Curtis distance, **H**) of the gut microbiota. **(H)** KEGG pathway enrichment analysis showing significant upregulation of butyrate metabolism pathways following VNS. **(I, M)** OPLS-DA score plots of untargeted metabolomics for feces (**I**) and plasma (**M**), showing distinct separation between Pre-VNS and Post-VNS samples. **(J, N)** Venn diagrams illustrating shared and unique differential metabolites in feces (**J**) and plasma (**N**) across time points. **(K, O)** Longitudinal changes in specific SCFAs. Fecal butyrate decreased (**K**), while plasma butyrate significantly increased (**O**) post-VNS. **(L, P)** Correlation between butyrate levels and clinical response. Fecal butyrate reduction was driven primarily by responders (R, n=6) compared to non-responders (N, n=5). Plasma butyrate elevation was driven primarily by responders compared to non-responders. **(Q)** Linear regression analysis revealed a significant positive correlation between the reduction in fecal butyrate at 6 months post-surgery and the reduction in seizure frequency; however, no significant correlation was found between the increase in plasma butyrate and the reduction in seizure frequency. **(R)** Linear regression analysis revealed a significant positive correlation between the reduction in fecal butyrate at 12 months post-surgery and the reduction in seizure frequency; however, no significant correlation was found between the increase in plasma butyrate and the reduction in seizure frequency. Data are presented as mean ± standard error, and individual data points are shown (E, F, G, K, L, O, P, Q and R). Paired t-tests were used (E, F, G, K, L, O, and P). Linear regression analysis (Q and R). *p < 0.05, **p < 0.01, ***p < 0.001, ns, no significant difference. VNS, Vagus nerve stimulation; pre, preoperatively; post 6M, six months postoperatively; post 12M, twelve months postoperatively; R, Response; N, Non-response; PCoA, principal coordinate analysis; OPLS-DA, Orthogonal partial least squares-discriminant analysis.

Metagenomic sequencing identified 2,220 genera and 10,496 species. Alpha diversity (Chao1 index) did not change at 6 months but decreased significantly by 12 months post-VNS (Figure 1F). Beta diversity (principal coordinate analysis, PCoA) showed no significant separation between preoperative and postoperative samples (adonis P=0.406; Figure 1G). LEfSe analysis (LDA >3.0, p<0.05) identified 19 differentially abundant taxa at 6 months and 22 taxa at 12 months compared to baseline (Figure 1D–E). Taxa enriched at 6 months included AKK (Figure 1F), *Prevotella*, *Alistipes*, and *Odoribacter*; at 12 months, Prevotella remained elevated. These taxa are established SCFAs producers(Gentile et al., 2023; Xu et al., 2021; Zhang et al., 2025). Consistent with this, KEGG enrichment analysis of metagenomic data revealed significant upregulation of butyrate metabolism post-VNS (Figure 1H).

Fecal untargeted metabolomics (Orthogonal partial least squares-discriminant analysis, OPLS-DA) showed clear separation between preoperative and postoperative samples (6 months: predictive component 7.04%, orthogonal 11%; 12 months: predictive 7.83%, orthogonal 11.2%; Figure 1I). After filtering unidentifiable metabolites, 412 and 526 differentially abundant metabolites were detected at 6 and 12 months, respectively, with 89 shared between both comparisons (Figure 1J, Table S3). The first 20 differential metabolites at both time points included butyrate (Figure S1G). Fecal butyrate levels decreased significantly at both postoperative time points; propionate decreased only at 6 months and valerate only at 12 months (Figure 1K). Stratification by response showed that fecal butyrate declined significantly in responders but not in non-responders (Figure 1L).

In plasma, untargeted metabolomics also distinguished postoperative from preoperative samples (6 months: predictive component 6.16%, orthogonal 9.71%; 12 months: predictive 7.75%, orthogonal 9.03%; Figure 1M). A total of 251 (6 months) and 370 (12 months) differentially abundant metabolites were identified, with 92 common to both comparisons (Figure 1N, Table S3). Notably, plasma butyrate increased significantly at both time points (Figure 1O). To clarify the relationship between the change in butyrate and the reduction in seizure frequency, a linear regression analysis was conducted to examine the relationship between the reduction in fecal butyrate and the reduction in seizure frequency, as well as the relationship between the increase in plasma butyrate and the reduction in seizure frequency. At 6 months post-surgery, the extent of decrease in fecal butyric acid was positively correlated with the extent of reduction in epileptic seizures (R^2^ = 0.5516, p = 0.0088, Figure 1Q). A similar positive correlation was observed at 12 months post-surgery (R^2^= 0.5806, p = 0.0064, Figure 1R). However, there was no significant correlation between the increase in plasma butyric acid and the reduction in epileptic seizures (Figure 1 Q-R).

Together, these data indicate that VNS remodels the gut microbiota in epilepsy patients, enriching SCFAs-producing bacteria such as AKK, *Prevotella*, *Alistipes*, and *Odoribacter*, and alters butyrate levels. The alteration of butyrate levels is significantly associated with clinical efficacy.

### VNS-remodeled microbiota attenuates seizure susceptibility and anxiety

We hypothesized that VNS exerts neuroprotective effects by reshaping the gut ecosystem. To uncouple the direct effects of stimulation from microbiota-mediated effects, we transplanted fecal microbiota from healthy VNS-treated donors into antibiotic-treated recipients prior to kainic acid (KA)-induced seizure induction (Figure 2A). VNS-conditioned microbiota was sufficient to lower seizure susceptibility; the FMT-VNS group demonstrated fewer and shorter seizure episodes than the FMT-sham controls (Figure 2B). Furthermore, FMT-VNS recipients exhibited improved emotional regulation in the open field test (OFT), showing increased locomotor activity and central zone preference (Figure 2C, Figure S2D). These results suggest that VNS induces a specific microbial landscape that prophylactically suppresses seizures and alleviates anxiety.

**Figure 2.**
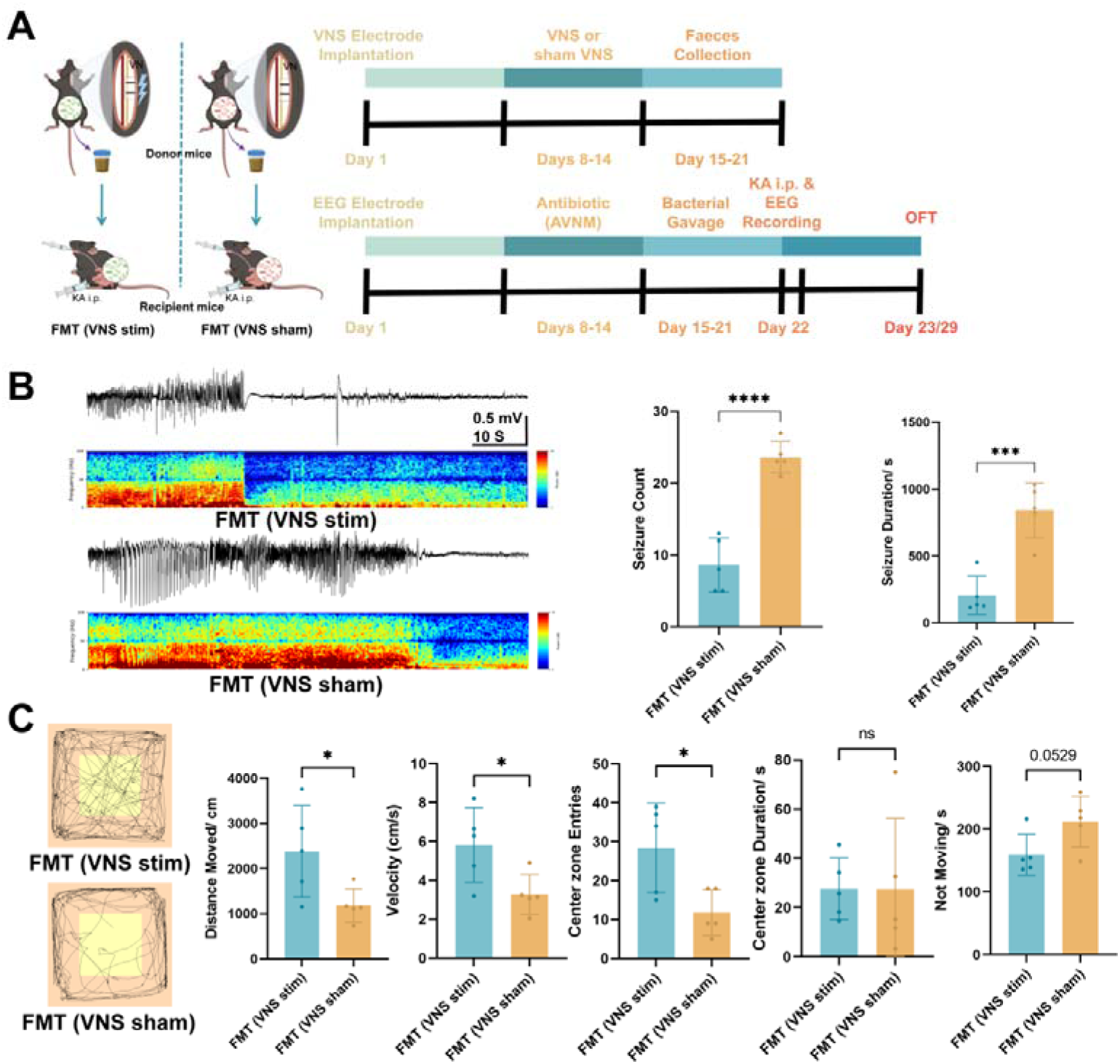
VNS-Conditioned Microbiota Attenuates Seizure Susceptibility in Recipient Mice. **(A)** Experimental schematic. Antibiotic-treated recipient mice FMT from donors subjected to active VNS (**FMT-VNS**, n=5) or sham stimulation (**FMT-Sham**, n=5) prior to KA-induced seizure modeling. **(B)** Representative EEG traces (left) and quantification of seizure parameters (right). The FMT-VNS group exhibited significantly reduced seizure frequency and shorter seizure duration compared to the FMT-Sham group. **(C)** Representative track plots from the OFT 1 day post-induction. Recipients of VNS-conditioned microbiota displayed increased total distance traveled, velocity, and center zone entries, along with reduced immobility time relative to the FMT-Sham group. Data are represented as mean ± SEM with individual data points shown (B and C). Statistical significance was determined by unpaired t-test. *p < 0.05, **p < 0.01, ***p < 0.001, ****p < 0.0001; ns, not significant. **FMT**, Fecal microbiota transplantation; **KA**, Kainic acid; **i.p.**, Intraperitoneal; **EEG**, Electroencephalogram; **AVNM**, Antibiotic cocktail (ampicillin, vancomycin, neomycin, and metronidazole); **OFT**, Open field test; **VNS**, Vagus nerve stimulation.

Previous studies have demonstrated that the gut microbiota of patients with epilepsy undergoes specific changes(Dahlin & Prast-Nielsen, 2019; Erickson & Barker-Haliski, 2024; Kundu et al., 2023; Yang et al., 2024). In a parallel experiment assessing the transmissibility of epileptic susceptibility, recipients of microbiota from KA-kindled donors (FMT-KA) developed more severe seizures and heightened anxiety-like behaviors compared to controls (Figure S2A–D). Thus, while epilepsy-derived microbiota exacerbates seizure severity, VNS-derived microbiota effectively counteracts these vulnerabilities.

### VNS increases gut *Akkermansia* abundance and modulates butyrate level in a murine epilepsy model

Given the multifactorial influences on human gut microbiota, we validated our clinical observations in a mouse model of epilepsy. Mice were implanted with vagus nerve electrodes and randomized to receive active VNS (VNS-stim) or sham stimulation (VNS-sham) after KA injection. Stimulation was applied 2□h/day for two weeks. Electroencephalogram (EEG) was recorded post-KA, and behavior was assessed in the open field after the stimulation period (Figure 3A).

**Figure 3.**
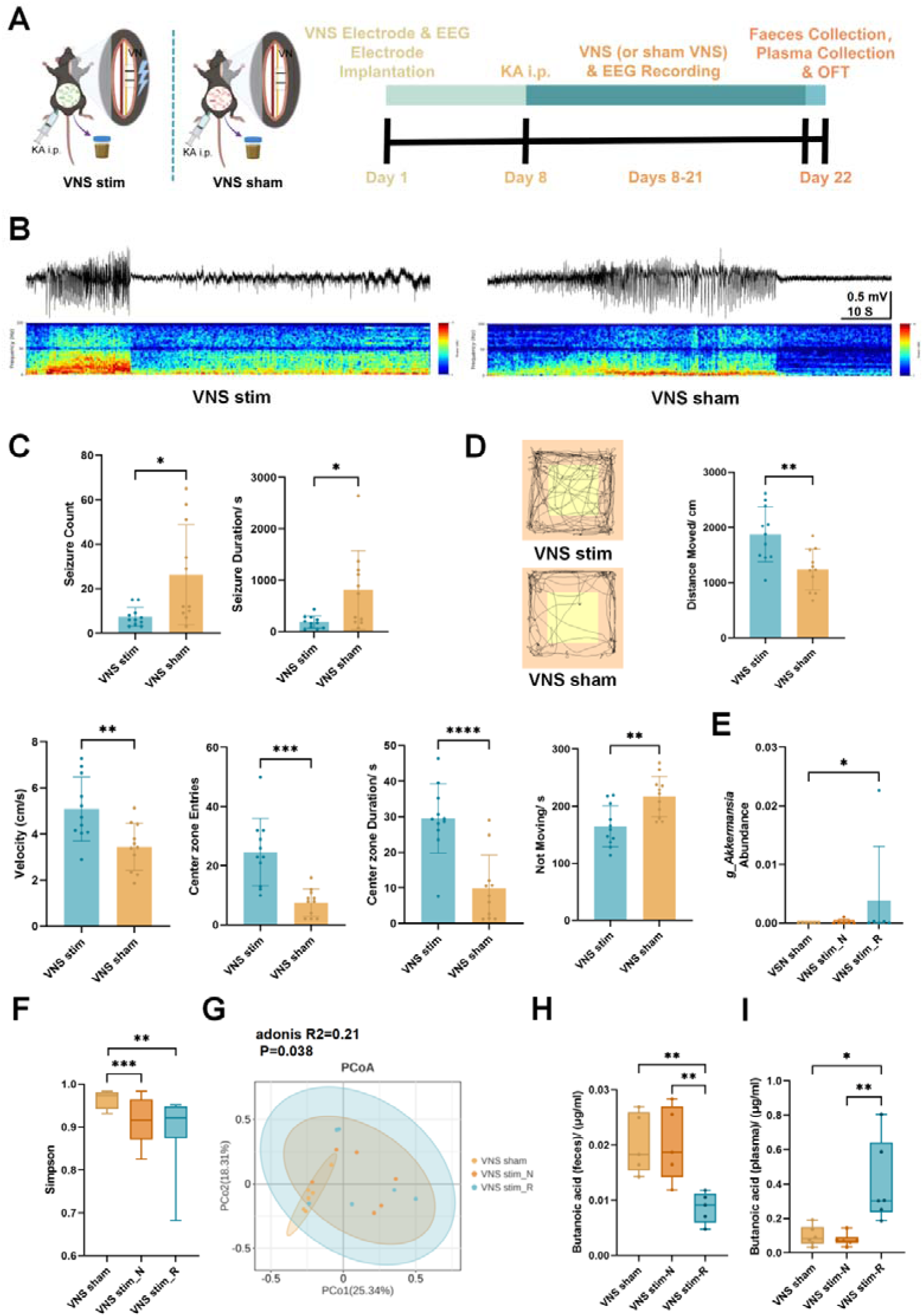
VNS Enriches *Akkermansia muciniphila* and Modulates Butyrate level in a Murine Model of Epilepsy. **(A)** Experimental design. Epileptic mice received either active VNS (VNS-stim, n=11) or sham stimulation (VNS-sham, n=11) for 2 weeks, followed by 16S rDNA sequencing and targeted SCFAs metabolomics. **(B, C)** VNS efficacy. Representative EEG traces (**B**) and quantification (**C**) confirming reduced seizure frequency and duration in the VNS-stim group. **(D)** Behavioral analysis. OFT trajectories and metrics showing reduced anxiety-like behavior in VNS-treated mice. **(E)** Relative abundance of *Akkermansia*. The enrichment of *Akkermansia* was specific to VNS responders (R, n=6) compared to non-responders (N, n=6) and sham controls. **(F, G)** Microbiota diversity analysis. Alpha diversity (Simpson index, **F**) and beta diversity (PCoA, **G**) indicate distinct community structures between groups. **(H)** Targeted metabolomics showing reduced fecal butyrate levels specifically in the VNS responder group. **(I)** Targeted metabolomics showing elevated plasma butyrate levels specifically in the VNS responder group. Data are presented as mean ± standard error, and individual data points are shown (C, D, E, H, and I). Unpaired t-tests were used (C, D, E, F, H, and I). *p < 0.05, **p < 0.01, ***p < 0.001, ****p < 0.0001. **VNS stim**, Vagus nerve real stimulation; **VNS sham**, Vagus nerve sham stimulation; **VN**, Vagus nerve; **KA**, Kainic acid; **i.p.**, Intraperitoneal injection; **EEG**, Electroencephalogram; **OFT**, Open field test; **R**, Response; **N**, Non-response; **PCoA**, Principal Co-ordinates Analysis.

VNS significantly reduced seizure frequency and duration compared to sham treatment (Figure 3B–C) and improved anxiety-like behaviors, as shown by increased locomotion, center-zone activity, and reduced immobility (Figure 3D).

Analysis of fecal microbiota (16S□rDNA sequencing) and plasma SCFAs (targeted metabolomics) revealed that VNS altered community structure. Alpha diversity (Shannon index) was lower in VNS-stim mice than in VNS-sham mice (Figure 3F), and beta diversity (PCoA) differed significantly between groups (adonis P=0.038; Figure 3G). When VNS-stim mice were subdivided by treatment response (responders, <10 seizures; non-responders, ≥10 seizures), responders exhibited higher relative abundance of *Akkermansia* than either non-responders or sham controls (Figure 3E). Consistent with the results in humans, in the respondents, the fecal butyrate level decreased (Figure 3H), while the plasma butyrate level increased (Figure 3I).

These controlled animal data reinforce the clinical findings, demonstrating that VNS enriches *Akkermansia* and alters butyrate levels in association with reduced seizure susceptibility.

### Gut microbiota from VNS-treated patients reduces seizure susceptibility in recipient mice

To determine whether VNS-remodeled gut microbiota confers protection against epilepsy, we performed FMT in mice. Recipient animals received fecal samples from the same epilepsy patients collected either before (VNS pre group) or 6 months after (VNS post group) VNS implantation. Prior to FMT, these mice underwent a 1-week antibiotic treatment. Following a 1-week FMT, mice were subjected to KA-induced seizures with simultaneous EEG recording. Anxiety-like behavior was assessed in the OFT at 1□day and 1□week post-KA (Figure 4A).

**Figure 4.**
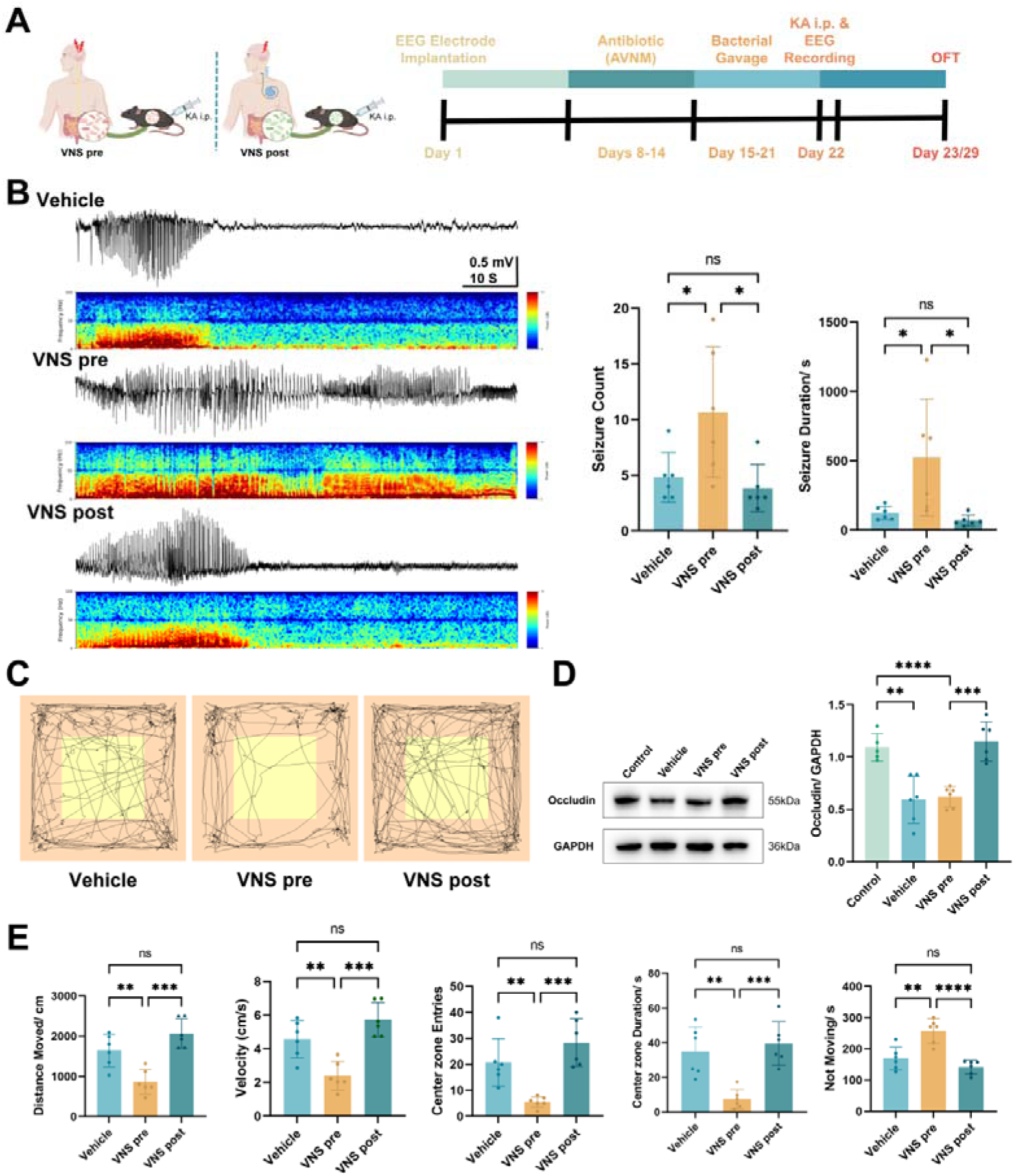
Fecal Microbiota Transplantation from VNS-Treated Patients Reduces Seizure Susceptibility and Restores Gut Barrier Integrity in Mice. **(A)** Schematic of the fecal microbiota transplantation (FMT) experiment. Antibiotic-treated mice received microbiota from patients collected either before VNS (VNS-pre, n=6) or 6 months post-VNS (VNS-post, n=6), or vehicle (PBS). **(B)** Representative EEG traces of kainic acid (KA)-induced seizures. Seizure quantification. Mice receiving VNS-post microbiota exhibited significantly fewer seizures (left) and shorter seizure duration (right) compared to the VNS-pre group. **(C, E)** Assessment of anxiety-like behavior in the OFT. Representative trajectories (**C**) and quantification (**E**) show increased locomotion and center zone exploration in the VNS-post group. **(D)** Western blot analysis of colonic Occludin expression. VNS-post microbiota transplantation restored Occludin levels that were downregulated in the VNS-pre group. Data are presented as mean ± standard error, and individual data points are shown (B, D, and E). Unpaired t-tests were used (B, D, and E). *p < 0.05, **p < 0.01, ***p < 0.001, ****p < 0.0001, ns, no significant difference. **VNS**, Vagus nerve stimulation; **pre**, preoperatively; **post**, postoperatively; **KA**, Kainic acid; **i.p.**, Intraperitoneal injection; **EEG**, Electroencephalogram; **AVNM**, ampicillin, vancomycin, neomycin and metronidazole; **OFT**, Open field test.

Mice receiving post-VNS microbiota exhibited significantly fewer seizures and shorter seizure duration compared to those receiving pre-VNS microbiota (Figure 4B), indicating that the gut microbial community after VNS treatment reduces seizure susceptibility. Notably, mice transplanted with pre-VNS microbiota showed higher seizure frequency and duration than vehicle-treated controls (Figure 4B), suggesting that the baseline microbiota of epilepsy patients may enhance seizure propensity.

In the OFT, the VNS post group displayed increased total distance traveled, higher velocity, more center-zone entries, longer center-zone duration, and reduced immobility at both time points compared to the VNS pre group (Figure 4C, E; Figure□S3A–B), indicating that post-VNS microbiota also improves affective behaviors with sustained effects.

We next examined whether post-VNS microbiota influences intestinal barrier integrity in epileptic mice. Western blot analysis of colon tissue revealed that Occludin—a key tight-junction protein—was markedly decreased in both vehicle and VNS pre groups relative to non-epileptic controls. In contrast, Occludin expression was significantly higher in the VNS post group than in the VNS pre group (Figure 4D), demonstrating that epilepsy-associated barrier impairment can be attenuated by VNS-modified microbiota.

Together, these findings show that the gut microbiota from VNS-treated epilepsy patients reduces seizure susceptibility, restores intestinal barrier function, and alleviates anxiety-like behavior in recipient mice.

### AKK reduces seizure susceptibility via the vagus nerve and modulates butyrate levels

Given the consistent enrichment of AKK following VNS in both patients and mice, we directly tested its anti-epileptic potential. Mice were pretreated with antibiotics and then gavaged with live AKK or vehicle for one week prior to KA challenge, with continuous EEG recording and subsequent open-field testing (Figure 5A).

**Figure 5.**
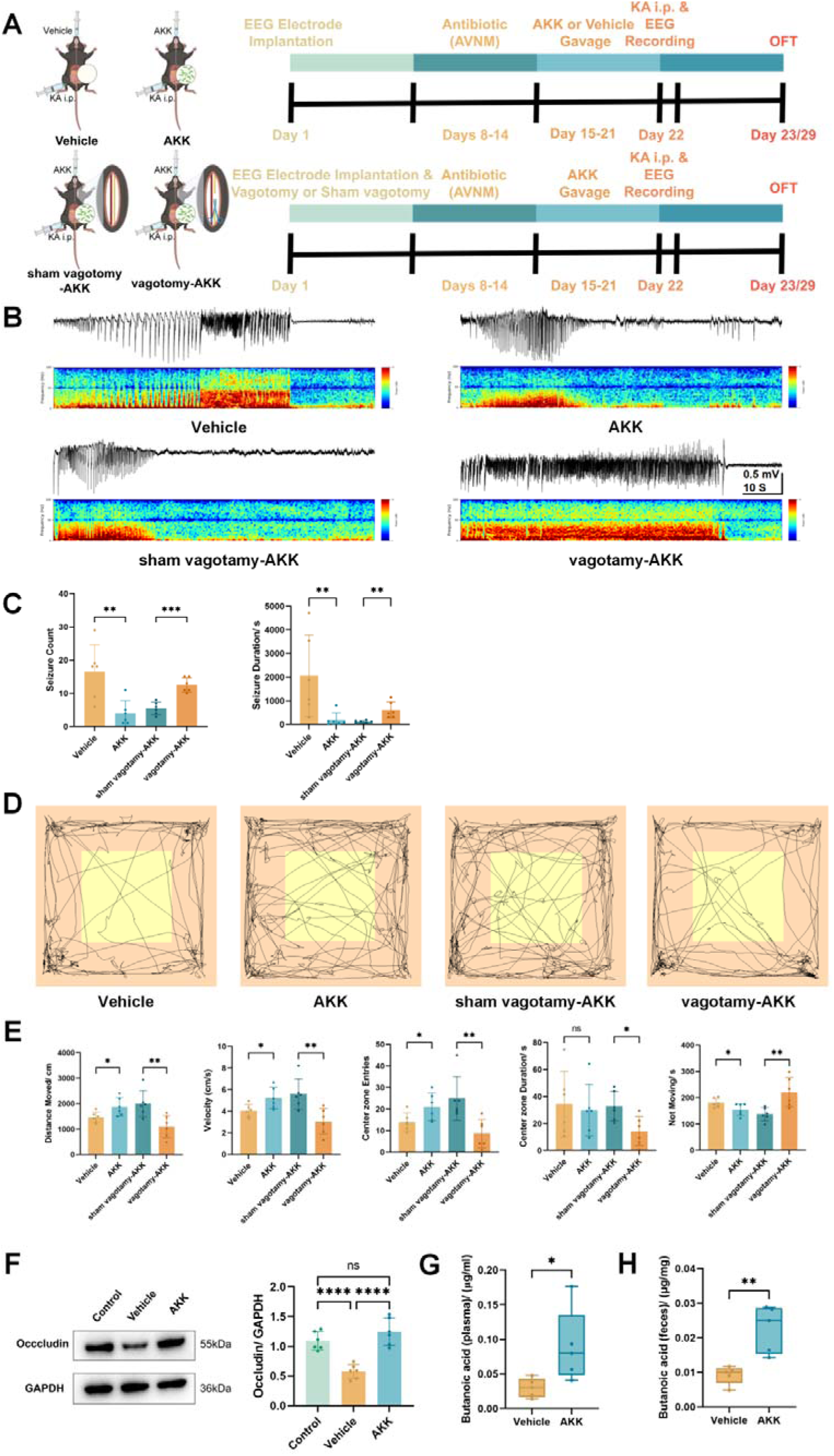
*Akkermansia muciniphila* Colonization Modulates butyrate levels and Attenuates Seizures via a Vagus Nerve-Dependent Pathway. **(A)** Experimental timeline. Mice were subjected to sham operation or subdiaphragmatic vagotomy prior to gavage with AKK or vehicle (n=6/group). **(B, C)** Seizure assessment. AKK colonization significantly reduced seizure frequency and duration compared to vehicle; this protective effect was abolished by vagotomy (**C**). **(D, E)** Anxiety-like behavior. The anxiolytic effects of AKK observed in the OFT were negated by vagotomy. **(F)** Colonic barrier function. AKK treatment upregulated Occludin protein expression compared to vehicle controls. **(G, H)** SCFAs levels. Targeted metabolomics confirmed that AKK colonization increased butyrate concentrations in both plasma (**G**) and feces (**H**). Data are presented as mean ± standard error, and individual data points are shown (C, E, F, G, and H). Unpaired t-tests were used (C, E, F, G, and H). *p < 0.05, **p < 0.01, ***p < 0.001, ****p < 0.0001, ns, no significant difference. **AKK**, *Akkermansia muciniphila*; **VN**, Vagus nerve; **KA**, Kainic acid; **i.p.**, Intraperitoneal injection; **EEG**, Electroencephalogram; **AVNM**, ampicillin, vancomycin, neomycin and metronidazole; **OFT**, Open field test.

AKK-colonized mice exhibited significantly fewer seizures and shorter seizure durations than vehicle-treated controls (Figure 5B–C). They also showed increased locomotion, greater center-zone exploration, and reduced immobility in the open field at both 1□day and 1□week post-KA (Figure 5D–E, Figure□S4A–B). Colon tissue analysis revealed higher Occludin expression in AKK-treated mice (Figure 5F), indicating improved intestinal barrier integrity.

To determine whether vagal signaling is required for AKK-mediated protection, we repeated the experiment in mice that underwent unilateral cervical vagotomy prior to AKK colonization. Vagotomy completely abolished the antiseizure and anxiolytic benefits of AKK (Figure 5B–E, Figure□S4A–B), confirming that AKK exerts its effects through a vagus nerve-dependent pathway.

Finally, we examined the link between AKK and butyrate, a key metabolite elevated after VNS. Targeted metabolomics showed that AKK colonization significantly increased butyrate levels in both feces and plasma compared to vehicle-treated mice (Figure 5G–H). Thus, AKK not only reduces seizure susceptibility and improves behavior in a vagally dependent manner, but also upregulates systemic butyrate, positioning it as a functional mediator within the VNS-driven microbiota–metabolite–brain axis.

### Butyrate reduces seizure susceptibility in mice through a vagus nerve-dependent mechanism

Given that VNS elevates systemic butyrate levels, we asked whether exogenous butyrate itself could attenuate epilepsy. Administration of butyrate (BA group) significantly reduced seizure frequency and duration compared to vehicle-treated controls (Figure 6B–C). butyrate also improved performance in the OFT, indicating alleviation of epilepsy-associated anxiety-like behavior, with effects persisting one week post-induction (Figure 6E–F, Figure□S5A–B). Moreover, colonic Occludin expression was higher in BA-treated mice, suggesting that butyrate helps restore intestinal barrier integrity compromised by epilepsy (Figure 6D).

**Figure 6.**
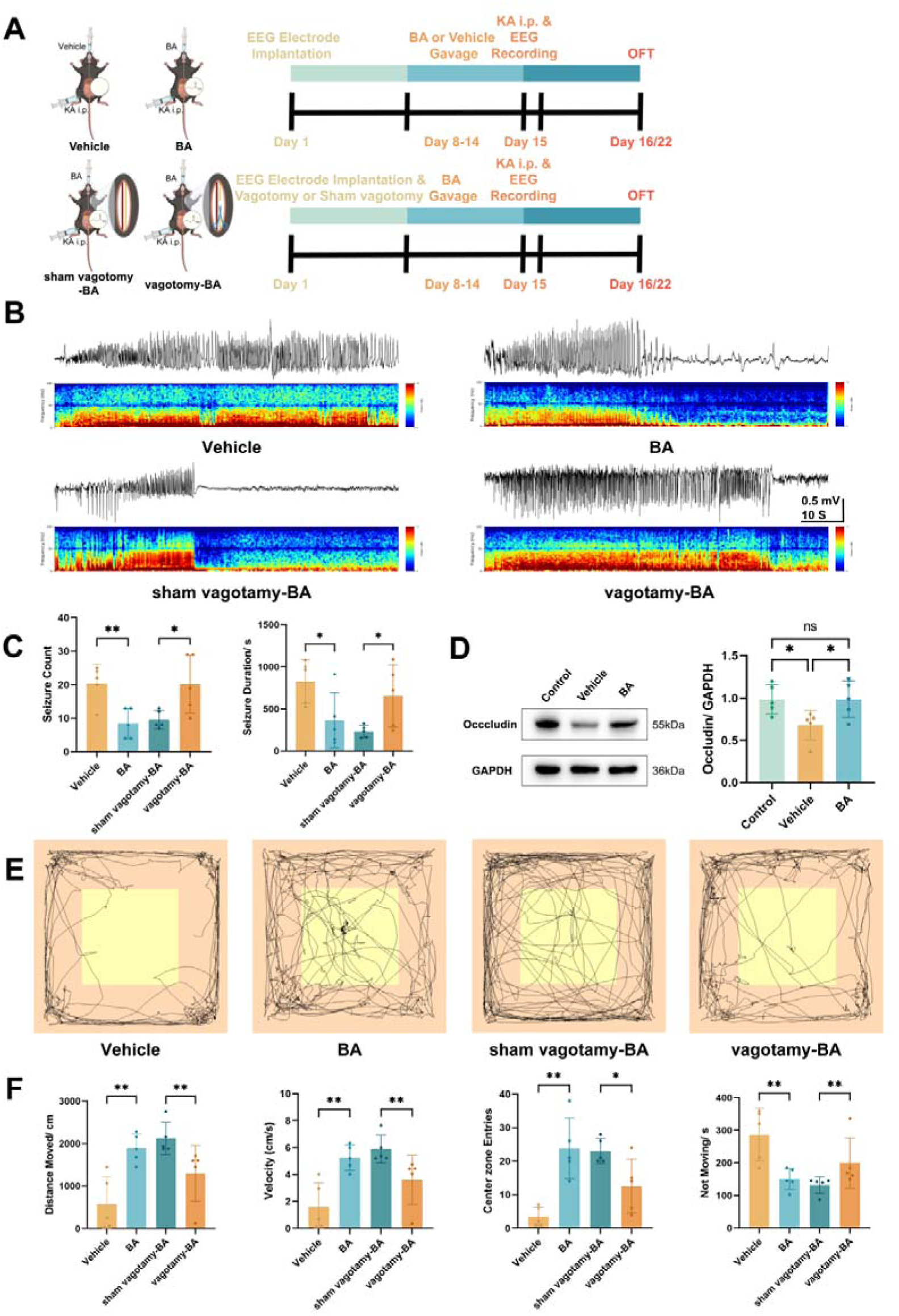
Exogenous Butyrate Mimics VNS Effects and Reduces Seizure Susceptibility in a Vagus Nerve-Dependent Manner. **(A)** Experimental design. Mice underwent sham operation or vagotomy followed by intragastric administration of butyrate (BA) or vehicle (PBS) for 1 week (n=5/group). **(B, C)** Seizure outcomes. Butyrate treatment significantly reduced seizure frequency and duration in sham-operated mice, but failed to protect vagotomized mice. **(D)** Gut barrier integrity. Butyrate administration restored colonic Occludin expression levels. **(E, F)** Behavioral analysis. The improvement in locomotor activity and anxiety-like behavior induced by butyrate was dependent on an intact vagus nerve. Data are presented as mean ± standard error, and individual data points are shown (C, D, and F). Unpaired t-tests were used (C, D, and F). *p < 0.05, **p < 0.01, ***p < 0.001, ****p < 0.0001, ns, no significant difference. **BA**, butyrate; **PBS**, phosphate buffer saline; **VN**, Vagus nerve; **KA**, Kainic acid; **i.p.**, Intraperitoneal injection; **EEG**, Electroencephalogram; **OFT**, Open field test.

To determine whether vagal signaling is required for these effects, we repeated the experiment in mice that underwent unilateral cervical vagotomy prior to butyrate administration. Vagotomy abolished all protective effects—seizure reduction, behavioral improvement, and barrier enhancement—observed in sham-operated animals (Figure 6C, E–F, Figure□S5A–B).

Thus, exogenous butyrate is sufficient to reduce seizure susceptibility and improve associated behavioral and intestinal deficits, and these benefits strictly depend on intact vagus nerve signaling.

## DISCUSSION

This study provides convergent clinical and preclinical evidence that VNS alleviates epilepsy by engaging a gut microbiota–butyrate–vagus nerve–brain axis. Our integrated approach, combining longitudinal patient profiling, multi-omics analyses, FMT, and targeted bacterial and metabolic interventions in mice, establishes a causal role for vagally mediated microbial remodeling in the antiseizure effects of VNS.

In patients with drug-resistant epilepsy, VNS not only reduced seizure frequency and severity but also consistently reshaped the gut microbiome, enriching taxa such as AKK(Dahlin & Prast-Nielsen, 2019; Erickson & Barker-Haliski, 2024; Kundu et al., 2023; Yang et al., 2024), *Prevotella*(Berni Canani et al., 2019; W. Lu et al., 2025), *Alistipes*(Chen et al., 2021; Shrode et al., 2023), and *Odoribacter*(Chen et al., 2021; Shrode et al., 2023), all implicated in SCFAs production. Metagenomic analysis revealed corresponding activation of butyrate metabolic pathways. Notably, fecal butyrate levels declined in clinical responders and correlated with improved outcomes, whereas plasma butyrate rose, suggesting altered host absorption or compartmentalization of microbial-derived butyrate after VNS. Parallel observations in a murine epilepsy model confirmed that VNS increased gut *Akkermansia* abundance, reduced fecal butyrate, and elevated plasma butyrate, further supporting the translatability of this microbial-metabolic shift.

We then demonstrated that AKK colonization alone was sufficient to modulate intestinal and plasma butyrate levels and reduce seizure susceptibility in mice. Although whether AKK directly produces butyrate or facilitates its production by other commensals remains to be fully resolved, our findings position this bacterium as a key mediator of VNS-driven metabolic and neurological benefits. Butyrate itself replicated the antiseizure and anxiolytic effects of VNS, and both bacterial and metabolic interventions required intact vagal signaling, highlighting the necessity of gut–brain communication via the vagus nerve.

Intriguingly, we observed an inverse relationship between fecal and plasma butyrate concentrations in both VNS-treated patients and mouse models. Numerous foundational studies have demonstrated that intestinal SCFAs, including butyrate, can signal to the brainstem either by acting on enterochromaffin cells or by directly stimulating receptors on vagal nerve endings, thereby modulating appetite, inflammation, and mood(Wu et al., 2021; Baslam et al., 2023; Mayer et al., 2023; Upreti et al., 2023). While previous studies confirm that butyrate can activate vagal signaling through receptors such as GPR43, GPR41, and Olfr78(Liu et al., 2024; Feng & Xu, 2023; Morales Fénero et al., 2021), our data suggest a specific kinetic mechanism. We hypothesize that VNS promotes the “local consumption” and “redistribution” of butyrate, facilitating its transport from the gut lumen into the bloodstream and enhancing its binding to vagus nerve-related receptors to further activate vagal signaling. This proposed mechanism of enhanced bioavailability warrants further investigation.

Butyrate, a well-characterized SCFA, exerts pleiotropic effects including immune modulation, enhancement of intestinal barrier function, and direct neuroactivity(Hill et al., 2021; Nugent et al., 2021; Zhou et al., 2025). Its increase post-VNS may suppress seizures through mechanisms involving oxidative stress reduction, mitochondrial stabilization, and neuronal excitability modulation. The concurrent enrichment of AKK, a next-generation probiotic linked to metabolic health, barrier integrity, and immunoregulation, suggests a synergistic role in maintaining gut–brain homeostasis(Ahmadi Badi et al., 2025; Ma et al., 2025b; Mu et al., 2025). Our work provides the first direct evidence linking AKK to seizure protection, extending its potential therapeutic relevance to epilepsy.

Notably, FMT from VNS-treated donors transferred seizure-protective and anxiolytic phenotypes to recipient mice, confirming that VNS-remodeled microbiota are functionally sufficient to confer therapeutic benefits. This underscores the gut microbiome not merely as a biomarker but as a modifiable effector in VNS therapy.

Several limitations of this study should be acknowledged. The clinical cohort was relatively small and lacked a healthy control group, and although antiepileptic drug regimens were held stable, confounding by unmeasured factors cannot be excluded. The KA model, while mechanistically informative, does not fully recapitulate human epilepsy. The precise molecular pathways, including butyrate receptors, epigenetic regulators, and vagal afferent signaling details, merit further investigation(C. Liu et al., 2025). The impact of VNS on the transport and utilization of butyrate also awaits further investigation. Future studies using gnotobiotic models, vagus-specific neuronal mapping, and multi-omics integration across larger, phenotypically diverse cohorts will help clarify translational relevance and interindividual variability.

In summary, we delineate a previously unrecognized pathway through which VNS ameliorates epilepsy: by enriching SCFAs-producing gut bacteria, modulating butyrate level, and engaging vagal signaling to reduce neuronal hyperexcitability. These findings reposition the gut–brain axis as a fundamental component of VNS mechanism of action and open new avenues for microbiota-targeted adjuvant therapies in epilepsy.

## RESOURCE AVAILABILITY

### Lead contact

Requests for further information and resources should be directed to and will be fulfilled by the lead contact, Yanwu Guo.

### Materials availability

All data associated with this study are present in the paper or the supplemental information and can also be obtained from the corresponding authors upon request.

### Data and code availability

Raw sequence data have been deposited in XXX

This paper does not report original code.

Any additional information required to reanalyze the data reported in this paper is available from the lead contact upon request.

## ACKNOWLEDGMENTS

We thank Dr. Xiaolong He for helpful discussions and insightful comments. The image of the mechanism was created with BioRender.com. We would like to thank all staff at the Department of Functional Neurosurgery and the Clinical Biobank Center of Zhujiang Hospital, Southern Medical University, for their technical assistance. We are also deeply grateful to all the donors and their families who kindly donated their samples to this study.

## AUTHOR CONTRIBUTIONS

H.S. and Y.G. conceptualized and organized the study. H.S. and L.Z. designed experiments and prepared the manuscript. P.C. and Z.Z. provided expertise and participated in the design of human study. Y.X., J.K., Y.L., Q.X., Y.Y., J.Y., S.X., and T.X. collected the samples and clinical information. J.Z. conducted bioinformatics analysis. L.Z., H.S and Z.C. interpreted the data. L.Z., Z.C., M.X., Y.W., A.Z., and Y.H. mainly completedanimal experiments. Y.G. and J.Z. revised the manuscripts. All authors reviewed and approved the final manuscript.

## STAR□METHODS

### KEY RESOURCES TABLE

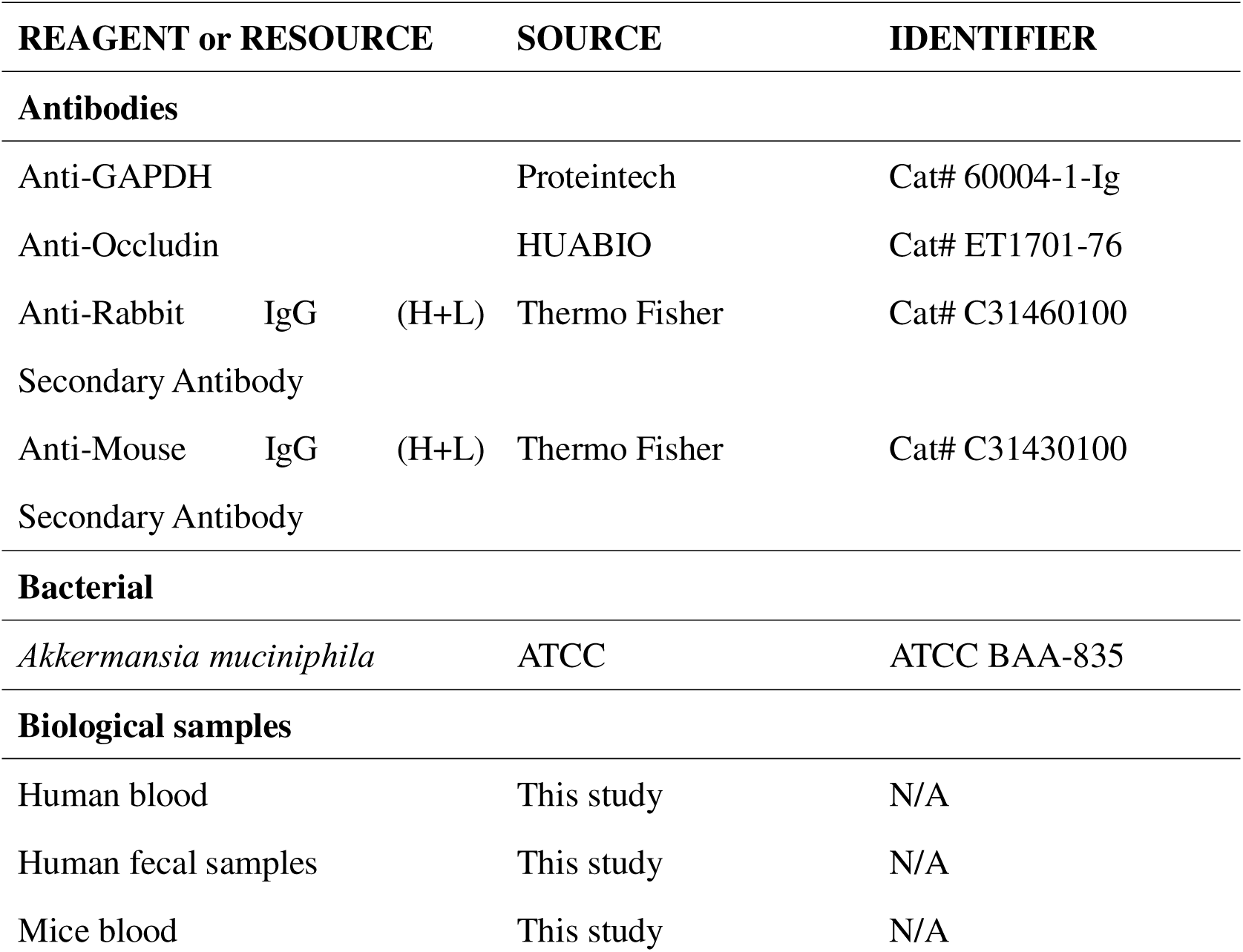
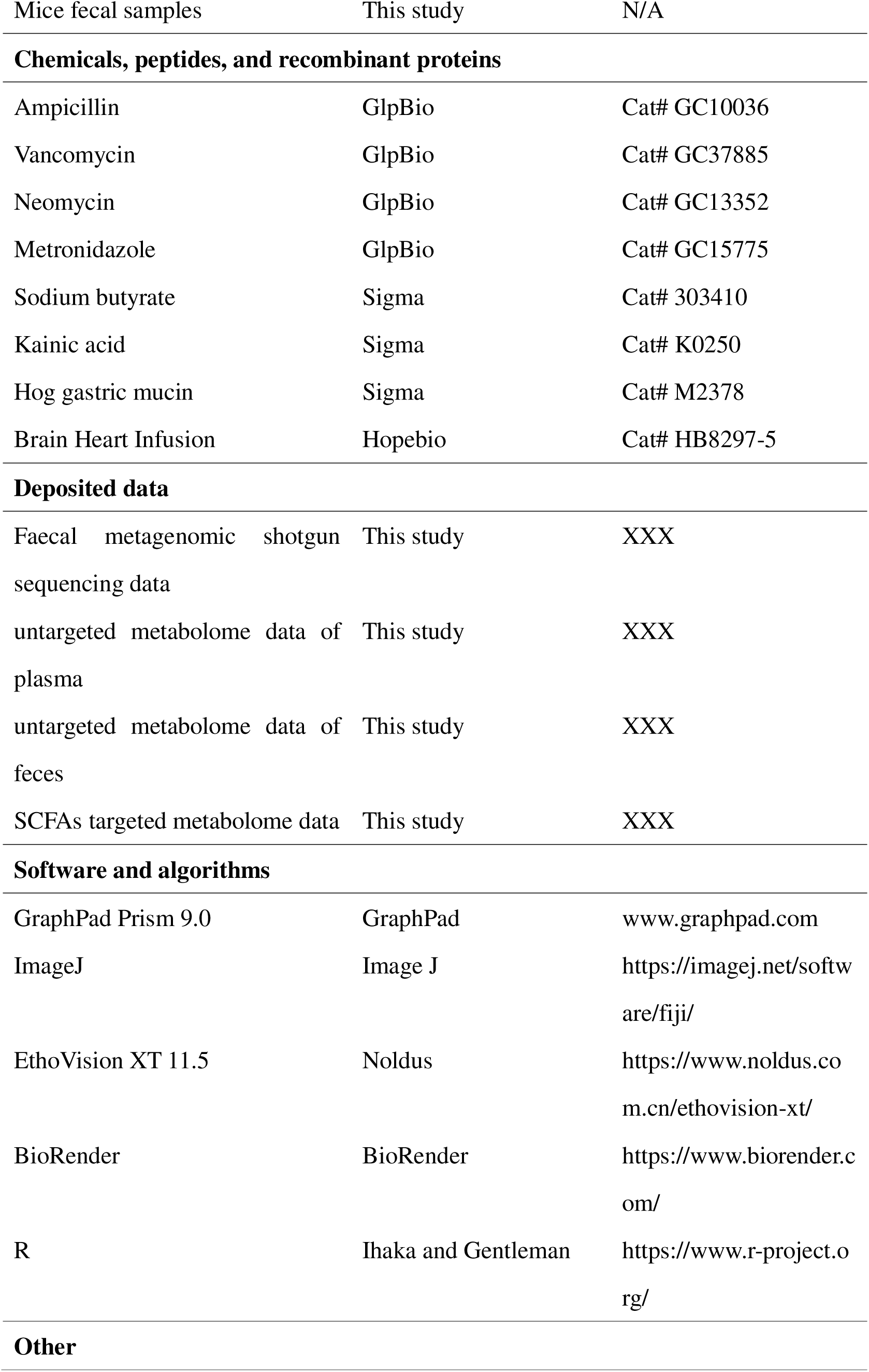
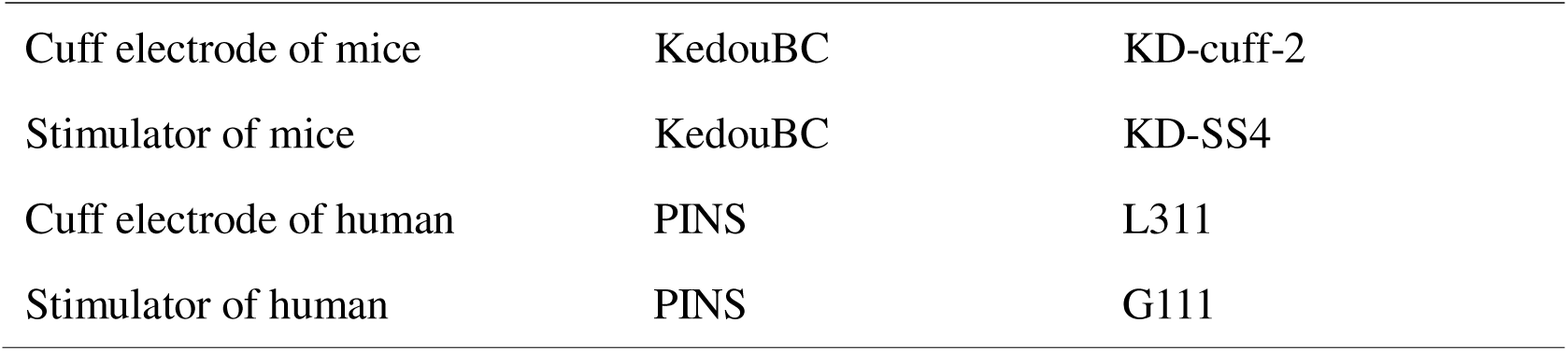

### Human study

This study was approved by the Ethics Committee of Zhujiang Hospital, Southern Medical University (2022-KY-096-02) and registered on the Chinese Clinical Trial Registry (ChiCTR2200055717). All patients or their guardians have given informed consent. The human study was designed as a single-center, single-arm clinical trial. As shown in Figure 1A, a total of 11 patients with drug-resistant epilepsy scheduled for vagus nerve stimulator implantation were enrolled sequentially based on strict inclusion and exclusion criteria and were followed up for one year. Three follow-up visits were arranged: preoperatively, 6 months postoperatively, and 12 months postoperatively. During each visit, plasma and stool samples were collected from the patients. Seizure conditions, including seizure frequency and the NHS3 score, were also assessed to evaluate the efficacy of VNS. Patients with a reduction in seizure frequency of 50% or more were classified as responders, while the others were classified as non-responders. Subsequently, the gut microbiome was analyzed using faecal metagenomic shotgun sequencing. Meanwhile, metabolomic analysis of stool and plasma samples was performed using liquid chromatography with tandem mass spectrometry (LC-MS/MS) technology. The vagus nerve stimulator was activated two weeks after surgery for all patients, and stimulation parameters were adjusted according to their clinical condition. Throughout the trial, the types of antiepileptic drugs remained unchanged, although doses could be adjusted for pediatric patients based on body weight changes. The inclusion and exclusion criteria are presented in Table S2.

### Patient sample collection

Fresh internal fecal samples were collected and immediately frozen at -80°C for subsequent use. Blood was collected in the morning using EDTA anticoagulant tubes after an overnight fast. The blood samples were then centrifuged at 1000×g and 4°C for 10 minutes to collect the plasma.

### Faecal metagenomic shotgun sequencing

The fecal samples used for metagenomic sequencing were extracted using the sodium dodecyl sulphate (SDS) method. DNA was subsequently diluted to 1ng/μL using sterile ddH2O, and its degradation degree and contamination were assessed on 1% agarose gels. DNA purity (OD260/OD280) was determined using the NanoDrop Microvolume Spectrophotometer (Thermo Fisher Scientific, USA). DNA concentration was measured using the Qubit® dsDNA Assay Kit in Qubit® 2.0 Fluorometer (Life Technologies, Carlsbad, CA, USA). All samples were sequenced on Illumina NovaSeq 6000 platform generating approximately 10 Gbp of 150-bp paired end reads per sample.

### Untargeted metabolite analysis

Untargeted metabolome measurements using ultra-high performance liquid chromatography - mass spectrometer (UHPLC-MS).

100 μL of plasma sample was transferred to an EP tube, followed by the addition of 300 μL of methanol containing 1 μg/mL L-2-chlorophenylalanine. The resulting mixture was vortexed for 30 seconds, sonicated for 10 minutes in an ice-water bath, and incubated for 1 h at -40 °C to precipitate proteins. Subsequently, the sample underwent centrifuged at 12000 rpm for 15 min at 4 °C. The resulting extract was then transferred to a clean glass vial for further analysis. A quality control (QC) sample was prepared by amalgamating equal aliquots of supernatants derived from each plasma sample(Dunn et al., 2011).

Weigh 25 mg of fecal sample at low temperature into an EP tube, add homogenizing beads, and then add 500 μL of extraction solution (methanol: acetonitrile: water = 2:2:1 (V/V)), which contains isotopically labeled internal standards. Vortex for 30 seconds. Homogenize in a homogenizer (35 Hz, 4 min), then transfer to an ice-water bath for ultrasonic treatment for 5 min. Repeat this step three times. Let it stand at -40°C for 1 hour. Centrifuge the sample at 4°C, 12000 rpm (centrifugal force 13800 (×g), radius 8.6 cm) for 15 min. Take the supernatant and put it into the injection bottle for machine detection. Take an equal amount of supernatant from all samples and mix them to form a QC sample for machine detection.

LC-MS/MS analyses were conducted utilizing a UHPLC 1290 system (Agilent Technologies, Waldbronn, Germany) coupled with a UPLC HSS T3 column (2.1 mm × 100 mm, 1.8 μm, Waters, Manchester, UK) and interfaced with a Q Exactive mass spectrometer (Orbitrap MS, Thermo Fisher Scientific, SanJose, CA). The extracts were gradient-eluted with water and acetonitrile with 0.1% formic acid for positive mode, or 5 mmol/L ammonium acetate for negative mode, and the mobile phase B was acetonitrile. The mass spectrometry was used for its ability to acquire MS/MS spectra on information-dependent acquisition (IDA) mode, controlled by Xcalibur 4.0.27 software (Thermo Fisher Scientific). The electrospray ionization (ESI) source conditions were configured as follows: sheath gas flow rate as 45 Arb, Aux gas flow rate as 15Arb, capillary temperature 400 °C, full MS resolution as 70000, MS/MS resolution as 17500, collision energy as 20/40/60 in NCE mode, spray Voltage as 4.0 kV (positive) or -3.6 kV (negative), respectively. To assess data repeatability, samples were randomized, and a QC sample was injected and analyzed after every 10 samples(Contrepois et al., 2020).

## Animal experimental protocol

### Mice

Male SPF C57BL/6 mice aged 6 to 8 weeks were housed in a standard laboratory environment with a temperature of 24 ± 1°C, humidity ranging from 50% to 70%, and a 12-hour light/12-hour dark cycle. They were provided with free access to food and water. The mice were fed a standard sterile diet and drinking water. All animal experiments were conducted in accordance with the guidelines for animal use, treatment, and euthanasia approved by the Animal Ethics Committee of Southern Medical University (License No. LAEC-2025-072).

### 16S rDNA gene sequencing

The fecal samples used for 16S *rDNA amplicon* metagenomic sequencing were extracted with the Power Soil DNA Isolation Kit (MO BIO Laboratories, Carlsbad, CA, USA) according to the manufacturer’s protocol. DNA quality and quantity were assessed by the ratios of 260 nm/280 nm and 260 nm/230 nm, respectively. Qualified DNA samples were amplified with 16S rDNA V3-V4 region using the forward primer (5′-ACTCCTACGGGAGGCAGCA-3′) and the reverse primer (5′-GGACTACHVGGGTWTCTAAT-3′) combined with adapter sequences and barcode sequences. Sequencing libraries were generated using TruSeq®DNA PCR-Free Sample Preparation Kit (Illumina, USA) following manufacturer’s recommendations and index codes were added. The library quality was assessed on the Qubit@ 2.0 Fluorometer (Thermo Scientific) and Agilent Bioanalyzer 2100 system. The sequencing was performed using the Illumina Novaseq 6000 (Illumina, San Diego, USA).

### Targeted metabolic analysis

SCFAs measurements using the gas chromatography - mass spectrometer (GC-MS). 50 μL plasma was mixed with 0.05 mL 50 % H2SO4 and 0.2 mL of 2-Methylvaleric acid (25 mg/L stock in methyl tert-butyl ether) as internal standard. The amalgamation was subjected to vortex mixing for 30 s, oscillations in 10 min, then ultrasound treated for 10 min with incubation in ice water. Subsequently, centrifugation was conducted for 15 min at 10000rpm, 4 °C. Keep at −20 °C for 30 min, the organic phase was collected and analyzed using SHIMADZU GC2030-QP2020 NX gas chromatography-mass spectrometer (Shimadzu Corporation, Kyoto, Japan) equipped with a HP-FFAP capillary column (30m×250μm×0.25μm, Agilent Technologies, Wilmington, DE, USA). A 1 μL aliquot of the analyte was injected in split mode (5:1). Helium was used as the carrier gas, with a front inlet purge flow of 3 mL min−1, and a gas flow rate through the column was 1 mL min−1. The initial temperature was maintained at 80 °C for 1 min, subsequently elevated to 200 °C at a rate of 10 °C min−1 for 5 min, and then sustained for 1 min at 240 °C at a rate of 40 °C min−1. The injection, transfer line, quad and ion source temperatures were 240 °C, 240 °C, 200 °C and 150 °C, respectively. The energy was -70 eV in electron impact mode. The mass spectrometry data were acquired in Scan/SIM mode with the m/z range of 33-150 after a solvent delay of 3.5 min. The RSD for internal standard in the QC samples averaged at 6.24% during the analysis of human plasma samples.

### Western blot analysis

After anesthesia with tribromoethanol (300 mg/kg) in the abdomen, the colon of mice was rapidly removed. Total protein of the tissue was extracted with lysis buffer (Cat# P0013B, Beyotime, Shanghai, China) and boiled in protein loading buffer. Equal amounts of denatured protein samples were electrophoresed in 6–10% polyacrylamide gels containing 0.1% SDS and transferred to polyvinylidene fluoride (PVDF) membranes with a pore size of 0.45 μm (Cat# IPVH00010, Millipore). When transferring proteins from the polyacrylamide gel to the PVDF membrane, the polyacrylamide gel, PVDF membrane and filter paper were stacked together and placed in a transfer sandwich. Then the sandwich was placed in a transfer tank and proteins in the polyacrylamide gel were transferred to the membrane at 250 mA resistance for 2 hours. The PVDF membrane was incubated with the primary antibody at 4°C for at least 12 hours, followed by incubation with the secondary antibody at room temperature for about 2 hours (Cat# C31460100, Thermo Fisher, USA; Cat# C31430100, Thermo Fisher, USA). Equal amounts of Clarity Western Peroxide reagent and Clarity Western Luminol/enhancer reagent (Cat# 1705060, Biorad, California, USA) were mixed together. The mixed reagent was added to the membrane and the required signal was visualized by a quantitative FluorChem SP imaging system (Alpha Innotech, California, USA). The integrated density of each band was measured by ImageJ software and the corresponding Gapdh band value was used as an internal control. Then the integrated density of Gapdh in the control group mice was normalized to 1 and the integrated densities of other bands were compared with Gapdh. The following primary antibodies were used: Occludin (1:500, Cat# ET1701-76, HUABIO, Hangzhou, China) and Gapdh (1:5000, Cat# 60004-1-Ig, Proteintech, Wuhan, China).

### OFT

Mice were acclimatized to the testing room for 1 h prior to the experiment. The test was conducted in a square open-field arena (Dimensions, e.g., 33.5 × 33.5 × 33.5 cm). Each mouse was gently placed in the center of the arena and allowed to explore freely for 6 min. Between trials, the apparatus was cleaned with 75% ethanol to remove olfactory cues. The distance moved, velocity, center zone entries, center zone duration, and not moving time were automatically recorded and analyzed using (EthoVision XT 11.5, Noldus).

### Vagotomy

Mice were anesthetized with isoflurane (flow rate: 1 L/min) and placed in a supine position on a surgical table after loss of consciousness. Following disinfection, a midline incision was made along the neck to expose the underlying muscle layer. The neck muscles were carefully dissected to locate the left carotid sheath. Under microscopic guidance, the vagus nerve was gently separated from the surrounding connective tissues while avoiding damage to the carotid artery and jugular vein. The nerve was then transected at an appropriate site to ensure complete disruption. The skin incision was sutured, and the animals were allowed to recover for one week(Zou et al., 2024).

### Epilepsy model establishment

A 10-minute baseline EEG recording (Solar, 1848) was performed. To investigate the susceptibility of mice to epilepsy, mice were administered a single intraperitoneal injection of KA (10 mg/kg, Cat# K0250, Sigma, USA). Immediately following the injection, EEG monitoring was continued for a duration of 2 hours.

### Vagus nerve stimulation

The vagus nerve was secured in a cuff electrode (KD-cuff-2, KedouBC, Suzhou, China), a bipolar device with the proximal lead serving as the cathode and the distal lead as the anode. The cuff electrode connector was implanted subcutaneously, while the stimulator connector was affixed to the skull using dental cement. The animals were allowed a 7-day postoperative recovery period. The VNS electrode was connected to a stimulatorr (KD-cuff-2, KedouBC, Suzhou, China). VNS was initiated immediately following KA injection, lasting 2 hours (parameters: frequency 30 Hz, pulse width 500 μs, current 1.5 mA, 30-second stimulation bursts with 5-minute inter-burst intervals). Subsequent stimulations were administered daily for 2 hours over a 2-week period.

### Antibiotic treatment

Vancomycin (50 mg/kg, Cat# GC37885, GlpBio, USA), neomycin (100 mg/kg, Cat# GC13352, GlpBio, USA), and metronidazole (100 mg/kg, Cat# GC15775, GlpBio, USA) were dissolved in sterile drinking water for mixing(Olson et al., 2018). The resulting mixture was then administered to mice via oral gavage at a frequency of twice daily for seven consecutive days. To ensure freshness and efficacy, the mixture was freshly prepared on a daily basis. Throughout the treatment period, ampicillin (1 mg/mL, GC10036, GlpBio, USA) was supplemented in the mice’s drinking water, with the solution being replaced with a fresh preparation every three days. All mice were maintained under sterile conditions.

### FMT

For FMT with mice as donors, fresh feces were collected from the disinfected anus of mice (10 mice per group) every day before the experiment to maintain the vitality of the microorganisms. The fresh feces were then weighed and mixed with sterile PBS at a dilution ratio of 1 mg/10 μL, and centrifuged at 900×g for 3 min. The supernatant was collected and administered by gavage to each mouse (10 mL/kg) for 7 days. All mice were treated under sterile conditions.

For human donor-derived FMT, frozen human fecal samples were thawed in a constant-temperature water bath at 37.5°C for 10 minutes. The remaining procedures were identical to those described previously. Unused supernatant was aliquoted and stored frozen. Repeated freeze-thaw cycles were strictly avoided.

### AKK transplantation

AKK (Cat# ATCC BAA845, ATCC, USA) was cultured under anaerobic conditions at 37°C in Brain Heart Infusion (Cat# HB8297-5, Hopebio, Qingdao, China) (media supplemented with hog gastric mucin (0.2%, Cat# M2378, Sigma, USA). 10□ CFU of bacteria were suspended in 200 μL of pre-reduced PBS and administered via oral gavage to mice that had undergone prior antibiotic treatment. The bacterial treatment was continued for one week.

### Butyrate treatment

Butyrate solution was freshly prepared in sterile PBS and administered to the mice via oral gavage at a dosage of 100 mg/kg body weight for 7 days (Cat# 303410, Sigma, USA)(Duan et al., 2025).

### EEG electrode implantation and EEG recording

Mice were anesthetized with isoflurane and secured in a stereotactic apparatus. Stainless steel microneedle electrodes for EEG recording were implanted into the skull above the hippocampus; the reference electrode was affixed within the skull over the prefrontal cortex, and the ground electrode was placed on the skull above the cerebellum. Finally, the electrodes were stabilized using dental adhesive. Subsequent experiments were initiated following a 1-week recovery period. Mouse EEGs were recorded using an EEG recording system (Solar, 1848) at a sampling rate of 1000 Hz. Data acquisition was performed with a band-pass filter set between 0.01 and 70 Hz. Seizure activity was assessed based on seizure frequency and duration.

### QUANTIFICATION AND STATISTICAL ANALYSIS

For each experiment, the number of independent biological replicates is denoted by n. The detailed information of n is provided in the legends accompanying each figure. The Shapiro-Wilk test was used to assess data normality, with p > 0.05 indicating that the data distribution did not significantly deviate from normality. If the data were normally distributed, a two-tailed Student’s t-test was used to compare two groups. A paired t-test was used to evaluate the changes before and after VNS treatment. One-way analysis of variance was used for comparisons across multiple groups. If the data were not normally distributed, non-parametric Mann–Whitney test (for two groups) or Kruskal–Wallis test (for multiple groups) was used for comparison. Data are presented as mean ± standard error of the mean (SEM), and a p-value < 0.05 was considered statistically significant. Specifically, *p < 0.05, **p < 0.01, ***p < 0.001, ****p < 0.0001. Detailed statistical methodology is provided in the legends accompanying each figure. All statistical analyses were performed using GraphPad Prism 9.0 or R. Statistical significance was set at p < 0.05.

