## Supplemental File for "Vagus Nerve Stimulation Ameliorates Epilepsy via a Gut Microbiota–Butyrate–Vagus Nerve–Brain Axis"

**Table S1. Patients’ demographics, baseline characteristics, and primary outcome results**

| **No.** | **Gender** | **Age at seizure onset**  **(y)** | **Age at initiation of  VNS (y)** | **Seizure types at initiation  of VNS** | **AEDs used at initiation of VNS** | **Seizure Frequency(**/ month**)**  **(pre)** | **Seizure Frequency(/** month**)**  **(post 6M)** | **Seizure Frequency (/** month**)**  **(post 12M)** | **NHS3**  **(pre)** | **NHS3**  **(post 6M)** | **NHS3**  **(post 12M)** |
| --- | --- | --- | --- | --- | --- | --- | --- | --- | --- | --- | --- |
| 1 | M | 4.2 | 4.7 | GTCS | OXC, VPA | 2 | 1 | 1 | 21 | 17 | 14 |
| 2 | M | 8.4 | 9.7 | GTCS | VPA, LEV, TPM, PHE, PIT | 30 | 20 | 3 | 17 | 15 | 14 |
| 3 | M | 0.3 | 4 | clonic | VPA, LEV | 3 | 2 | 2 | 11 | 11 | 11 |
| 4 | M | 1.2 | 4.2 | atonic | VPA, LTG,  PIT, VGB | 15 | 13 | 9 | 11 | 10 | 10 |
| 5 | M | 13.0 | 21.8 | GTCS | VPA, TPM | 1 | 0 | 0 | 12 | 1 | 1 |
| 6 | F | 1.1 | 9.2 | GTCS、tonic | OXC, TPM,  LTG, CZP | 50 | 42 | 40 | 21 | 21 | 19 |
| 7 | M | 2.2 | 10.5 | clonic | LEV, VPA, OXC, TPM, ZON | 45 | 45 | 40 | 10 | 10 | 10 |
| 8 | F | 10.2 | 14.5 | FS、tonic | LEV, TPM,  CZP, LAC | 92 | 44 | 21 | 13 | 11 | 11 |
| 9 | F | 2.2 | 3.9 | clonic、GTCS | VPA, TPM,  LEV, VGB | 180 | 162 | 157 | 10 | 10 | 10 |
| 10 | F | 16.0 | 32.5 | GTCS、absence | LEV, TPM | 1 | 0 | 0 | 12 | 1 | 1 |
| 11 | F | 3.3 | 4.5 | FS、clonic、tonic | VPA, LTG,  VGB, PHE | 450 | 150 | 135 | 14 | 12 | 10 |

Abbreviations: **y**, year; **VNS**, vagus nerve stimulation; **NHS3**, National Hospital Seizure Severity Scale; **pre**, preoperatively; **post 6M**, six months postoperatively; **post 12M**, twelve months postoperatively; **M**, Male; **F**, Female; **GTCS**, Generalized Tonic-Clonic Seizure; **FS**, Focal Seizure; **OXC**, Oxcarbazepine; **VPA**,Sodium Valproate; **LEV**,Levetiracetam; **TPM**,Topiramate; **PHE**,Phenobarbital; **PIT**,Pitolisant; **VGB**,Vigabatrin; **LTG**, Lamotrigine; **CZP**, Clonazepam; **ZON**, Zonisamide; **LAC**, Lacosamide

**Table S2. Inclusion and exclusion criteria for human trials**

| **No.** | **Inclusion criteria** |
| --- | --- |
| 1 | Patients diagnosed with drug-resistant epilepsy (DRE) by an epilepsy specialist, meeting the 2014 International League Against Epilepsy (ILAE) diagnostic criteria for epilepsy (citation). |
| 2 | Seizure frequency ≥1 episode per month. |
| 3 | No treatable cause of epilepsy identified, or failure of targeted etiology treatment. |
| 4 | Patients or their guardians fully understood the study and voluntarily participated. |
| 5 | Patients or their guardians demonstrated good compliance and were able to complete postoperative follow-up as required. |
| **No.** | **Exclusion criteria** |
| 1 | History of bilateral vagus nerve injury or transection. |
| 2 | Local infection at the neck or chest implantation site. |
| 3 | Intolerance to foreign object implantation or being prone to rejection. |
| 4 | Poor systemic condition unable to tolerate anesthesia and surgery. |
| 5 | Requirement for microwave or short-wave diathermy at the left chest or neck implantation site. |
| 6 | Presence of obstructive sleep apnea syndrome, severe digestive system diseases, hereditary metabolic disorders, severe cardiac conduction block. |
| 7 | Have used antibiotics, probiotics or prebiotics in the past three months. |

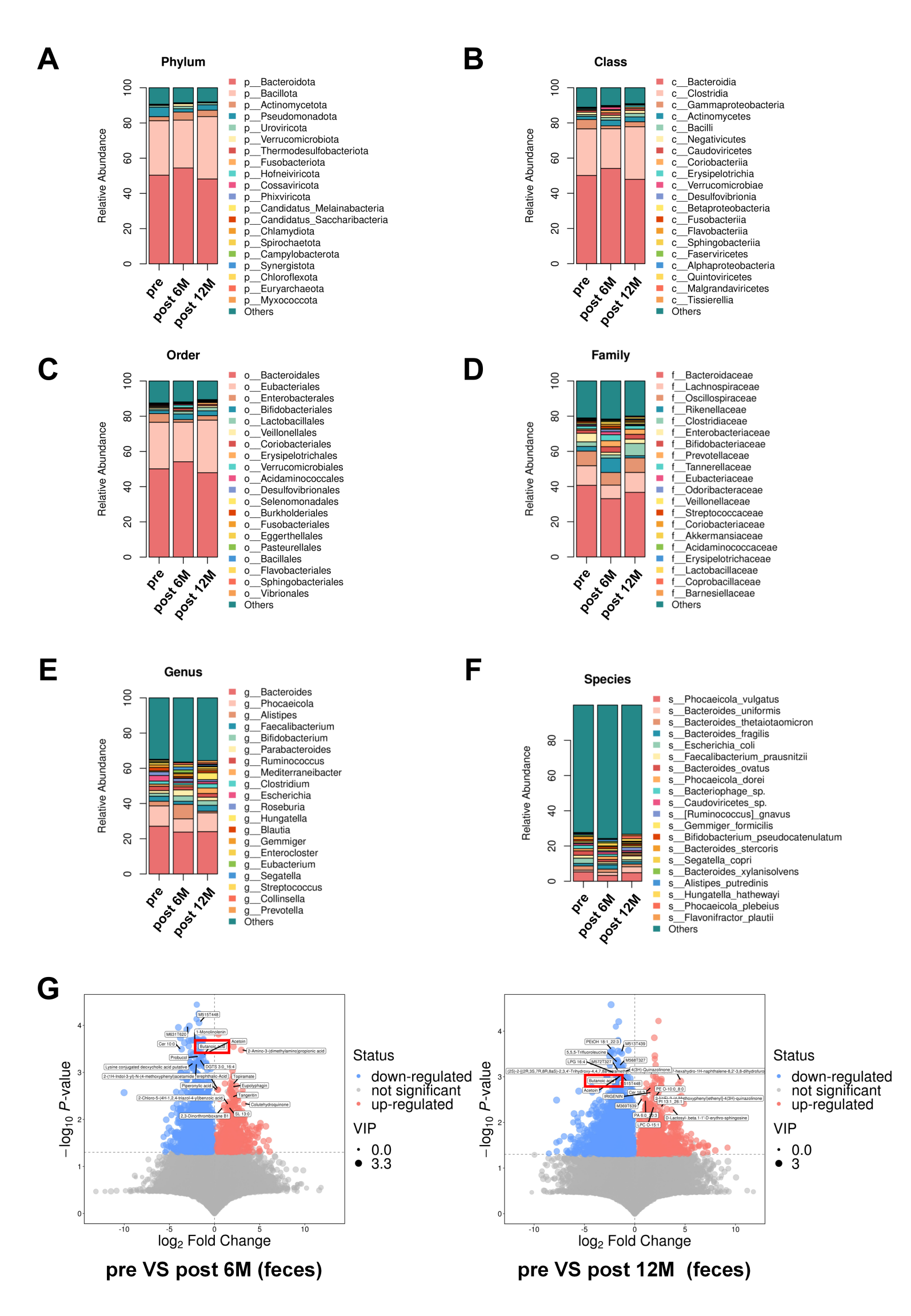

**Figure S1. Changes in the abundance of gut microbiota across hierarchical taxonomic levels in VNS patients (top 20 taxa).**

**(A)** At the phylum level, the relative abundances of the top 20 taxa across preoperative, 6 months postoperative, and 12 months postoperative show increases in *Actinomycetes*, *Bacilli*, and *Caudoviricetes*, and a decrease in *Betaproteobacteria*.

**(B)** At the class level, the relative abundances across timepoints show increases in *Actinomycetota* and *Uroviricota* and decreases in *Pseudomonadota* and *Fusobacteriota*.

**(C)** At the order level, the relative abundances across timepoints show increases in *Bifidobacteriales*, *Lactobacillales*, and *Coriobacteriales* and a decrease in *Enterobacterales*.

**(D)** At the family level, the relative abundances across timepoints show increases in *Bifidobacteriaceae*, *Prevotellacea*e, and *Tannerellaceae* and decreases in *Bacteroidaceae* and *Enterobacteriaceae*.

**(E)** At the genus level, the relative abundances across timepoints show increases in *Bifidobacterium* and *Parabacteroides* and decreases in *Bacteroides*, *Phocaeicola*, *Ruminococcus*, and *Escherichia*.

**(F)** At the species level, the relative abundances across timepoints show increases in *Bacteroides uniformis*, *Caudoviricetes sp.*, *Ruminococcus gnavus*, *Gemmiger formicilis*, and *Segatella copri* and decreases in *Phocaeicola vulgatus*, *Bacteroides thetaiotaomicron*, *Escherichia coli*, *Bacteroides ovatus*, *Bacteriophage sp.*, *Bacteroides stercoris*, and *Bacteroides xylanisolvens*.

**(G)** Volcano plots of differential fecal metabolites comparing preoperative vs 6 months postoperative and preoperative vs 12 months postoperative show a significant decrease in butyrate in both comparisons.

**Table S3. At two postoperative time points, the contents of 89 fecal metabolites all showed significant changes**

| **Metabolite** | **MEAN**  **(pre)** | **MEAN**  **(post 6 month)** | **MEAN**  **(post 12 month)** | **P-VALUE**  **(pre vs post 6 month)** | **P-VALUE**  **(pre vs post 12 month)** |
| --- | --- | --- | --- | --- | --- |
| cis-11.14-Eicosadienoic acid | 24.6297 | 13.1415 | 13.0294 | 0.0251 | 0.0167 |
| Butanoic acid | 267.9678 | 109.5943 | 106.9123 | 0.0004 | 0.0009 |
| Nervonic acid | 5.2117 | 2.3945 | 2.5380 | 0.0174 | 0.0223 |
| Terephthalic-Acid | 0.5157 | 0.6736 | 0.8143 | 0.0023 | 0.0100 |
| 2-Phenylpropionic acid | 32.7601 | 15.2379 | 10.6727 | 0.0409 | 0.0067 |
| 16-Hydroxypalmitic acid | 12.6315 | 6.7949 | 6.7453 | 0.0468 | 0.0401 |
| 2-Hydroxypalmitic acid | 12.6315 | 6.7949 | 6.7453 | 0.0468 | 0.0401 |
| Propionic acid | 12.6938 | 6.9029 | 8.6791 | 0.0007 | 0.0191 |
| Deoxycholic acid | 334.5450 | 211.6088 | 187.6119 | 0.0221 | 0.0106 |
| N-Acetylgalactosamine | 0.2718 | 0.0618 | 0.0712 | 0.0392 | 0.0475 |
| 4(3H)-Quinazolinone | 0.1006 | 0.0645 | 0.0504 | 0.0175 | 0.0007 |
| Chenodeoxycholic acid | 338.2073 | 213.9662 | 189.8038 | 0.0223 | 0.0107 |
| Indole-3-propionic acid | 0.0783 | 0.0251 | 0.0264 | 0.0109 | 0.0155 |
| DHA ethyl ester | 1.2970 | 0.7016 | 0.4066 | 0.0458 | 0.0062 |
| 3-Phenylpropanoic acid | 32.8421 | 15.2807 | 10.7026 | 0.0410 | 0.0067 |
| 4-Allylcatechol | 37.4986 | 16.3524 | 10.4113 | 0.0396 | 0.0071 |
| PC(18:1(9Z)/18:1(9Z)) | 0.4616 | 0.1467 | 0.2296 | 0.0041 | 0.0372 |
| PC(18:1(9E)/18:1(9E)) | 0.4532 | 0.1440 | 0.2254 | 0.0041 | 0.0372 |
| PC(18:1(6Z)/18:1(6Z)) | 0.4519 | 0.1436 | 0.2248 | 0.0041 | 0.0372 |
| 2'-Hydroxy-5'-methylacetophenone | 37.3151 | 16.2613 | 10.3533 | 0.0393 | 0.0070 |
| 4'-Hydroxy-3'-methylacetophenone | 37.4986 | 16.3524 | 10.4113 | 0.0396 | 0.0071 |
| 4′-Hydroxy-2′-methylacetophenone | 37.2431 | 16.2385 | 10.3388 | 0.0395 | 0.0071 |
| Acetoin | 265.1449 | 108.4399 | 105.8159 | 0.0004 | 0.0009 |
| 2'-Hydroxy-4'-methylacetophenone | 32.8792 | 15.2908 | 10.7097 | 0.0408 | 0.0067 |
| Piperonylic acid | 0.4750 | 0.6204 | 0.7501 | 0.0023 | 0.0100 |
| cis-4,10,13,16-Docosatetraenoic acid | 18.9904 | 6.3136 | 8.6290 | 0.0094 | 0.0277 |
| Cer 14:0 | 0.1195 | 0.0095 | 0.0209 | 0.0204 | 0.0351 |
| Stemphol | 0.1726 | 0.2294 | 0.2357 | 0.0329 | 0.0248 |
| Cecropiacic acid | 1.0057 | 0.5224 | 0.3163 | 0.0403 | 0.0055 |
| 2-(3-Methylbutoxy)acetic acid | 0.1444 | 0.0608 | 0.0401 | 0.0049 | 0.0010 |
| 8-iso-15-keto-PGE2 | 0.0152 | 0.4668 | 0.0787 | 0.0481 | 0.0138 |
| DG O-18:5_5:0 | 0.2309 | 0.1124 | 0.1225 | 0.0134 | 0.0335 |
| Probucol | 35.9441 | 11.0074 | 13.5678 | 0.0005 | 0.0011 |
| LPE O-16:1 | 6.1426 | 10.5968 | 12.7050 | 0.0338 | 0.0247 |
| MGDG O-11:0_2:0 | 7.9828 | 3.9513 | 4.2128 | 0.0318 | 0.0408 |
| 2,6-Dimethyl-4-hydroxybenzaldehyde | 37.3151 | 16.2613 | 10.3533 | 0.0393 | 0.0070 |
| Eicosapentaenoic acid methyl ester | 0.9851 | 0.1180 | 0.2229 | 0.0010 | 0.0024 |
| Lysine conjugated deoxycholic acid putative | 0.4160 | 0.0854 | 0.1877 | 0.0007 | 0.0229 |
| neobavaisoflavone | 0.2500 | 0.0571 | 0.0871 | 0.0086 | 0.0253 |
| 3H-Benzimidazol-5-yl-[4-[(5-cyclopropyl- 1H-pyrazol-3-yl)amino]pyrimidin-2-yl]amine | 0.2266 | 0.0056 | 0.0167 | 0.0344 | 0.0425 |
| 5.alpha.-Pregnane-3.beta.,17-diol-20-one | 3.0702 | 0.2283 | 0.8295 | 0.0010 | 0.0052 |
| Leucine conjugated chenodeoxycholic acid putative | 0.2476 | 0.0444 | 0.0440 | 0.0452 | 0.0451 |
| CAR 17:2 | 1.3883 | 0.6894 | 0.5343 | 0.0055 | 0.0009 |
| 3,12-dihydroxy-4,6a,6b,11,12,14b-hexamethyl -1,2,3,4a,5,6,7,8,9,10,11,12a,14,14a-tetradecahydropicene-4,8a-dicarboxylic acid | 1.7699 | 0.1829 | 0.4957 | 0.0102 | 0.0312 |
| 5-Bromo-6-nitro-1H-isoindole-1,3(2H)-dione | 0.0239 | 0.0109 | 0.0441 | 0.0032 | 0.0466 |
| 3-(2-Benzothiazolyl)coumarin | 0.2798 | 0.0422 | 0.0496 | 0.0290 | 0.0331 |
| PC(18:1(9Z)/18:0) | 2.7320 | 0.8503 | 0.6961 | 0.0132 | 0.0075 |
| VAE 18:4 | 1.6361 | 0.6466 | 0.8187 | 0.0099 | 0.0274 |
| Acetildenafil | 1.2949 | 0.5088 | 0.4519 | 0.0287 | 0.0198 |
| Oryzarol | 0.2169 | 0.1180 | 0.1039 | 0.0069 | 0.0072 |
| Methanone, 1,1'-[2-(phenylamino)-1,4- phenylene]bis[1-[4-(1-pyrrolidinyl)-1-piperidinyl]- | 0.8061 | 0.2097 | 0.1573 | 0.0356 | 0.0257 |
| MG(0:0/18:3(6Z,9Z,12Z)/0:0) | 0.8734 | 0.1031 | 0.2538 | 0.0011 | 0.0051 |
| N-Tetradecanoyl-DL-homoserine lactone | 1.4683 | 0.3429 | 0.3952 | 0.0285 | 0.0366 |
| Methyl 3-[(2-methyl-1,3-thiazolidin-4-yl) formamido]propanoate | 0.1099 | 0.0513 | 0.0655 | 0.0153 | 0.0361 |
| DGTS 4:0_13:1 | 2.1263 | 0.8242 | 0.6267 | 0.0462 | 0.0247 |
| DG 8:0_16:4 | 4.7765 | 1.8186 | 2.1834 | 0.0206 | 0.0464 |
| Jasminoside Q | 0.1318 | 0.0418 | 0.0417 | 0.0447 | 0.0415 |
| 1-Monolinolenin | 2.1213 | 0.4103 | 0.7565 | 0.0002 | 0.0018 |
| 9,11-Methane-epoxyprostaglandin F1.alpha. | 4.4688 | 0.2490 | 0.8306 | 0.0185 | 0.0374 |
| Cys-Ser | 0.1248 | 0.0548 | 0.0639 | 0.0058 | 0.0318 |
| .beta.-Hydroxyphenylalanine | 8.4927 | 1.8047 | 2.0195 | 0.0390 | 0.0449 |
| Gelomulide N | 0.1344 | 0.0300 | 0.0443 | 0.0206 | 0.0435 |
| FAHFA 4:0/18:2 | 0.4616 | 0.1059 | 0.1174 | 0.0015 | 0.0020 |
| Camylofine | 0.9886 | 0.4406 | 0.4422 | 0.0084 | 0.0096 |
| 2-Methoxy-4-((1R,2S,4R,6S)-5,5,6- trimethylbicyclo[2.2.1]heptan-2-yl)phenol | 0.3869 | 0.1872 | 0.1708 | 0.0135 | 0.0053 |
| 6-Ketotestosterone enanthate | 1.2765 | 0.7149 | 0.6993 | 0.0276 | 0.0303 |
| LPG 20:2 | 1.6775 | 0.5586 | 0.4253 | 0.0392 | 0.0271 |
| Goshonoside F2 | 28.8724 | 7.6960 | 10.6529 | 0.0124 | 0.0283 |
| Ethoxyquin | 0.4992 | 0.0873 | 0.1576 | 0.0168 | 0.0437 |
| PS(18:0/18:1(9Z)) | 1.4583 | 0.5746 | 0.5754 | 0.0134 | 0.0147 |
| L-Ala-Gly-Gly-Gly | 0.1799 | 0.0529 | 0.0569 | 0.0334 | 0.0372 |
| 2-Chloro-4-fluoro-5-nitrobenzoic acid | 0.2395 | 0.1486 | 0.5719 | 0.0136 | 0.0210 |
| M513T439 | 6.8463 | 3.4233 | 2.5146 | 0.0075 | 0.0003 |
| M481T482 | 555.4726 | 188.6075 | 254.8265 | 0.0119 | 0.0395 |
| M497T461 | 14.8241 | 4.9922 | 5.0233 | 0.0222 | 0.0221 |
| M515T417 | 4.5463 | 1.9820 | 1.7510 | 0.0091 | 0.0054 |
| M515T448 | 60.3183 | 18.4135 | 24.3020 | 0.0001 | 0.0009 |
| M499T477 | 21.8818 | 4.8592 | 6.6413 | 0.0125 | 0.0216 |
| M477T526 | 1.0952 | 0.2501 | 0.1310 | 0.0175 | 0.0087 |
| M485T411 | 2.7243 | 1.1023 | 0.8324 | 0.0402 | 0.0147 |
| M401T210 | 0.0880 | 0.0244 | 0.0289 | 0.0134 | 0.0211 |
| M572T327 | 0.1751 | 0.0841 | 0.0590 | 0.0225 | 0.0007 |
| M419T644 | 0.0346 | 0.0147 | 0.0192 | 0.0030 | 0.0230 |
| M484T443 | 5.8379 | 2.7279 | 2.4373 | 0.0289 | 0.0178 |
| M454T468 | 2.1622 | 1.3605 | 0.8640 | 0.0266 | 0.0041 |
| M397T431 | 0.4852 | 0.2902 | 0.2868 | 0.0428 | 0.0436 |
| M514T422 | 0.4196 | 0.0881 | 0.0597 | 0.0188 | 0.0122 |
| M568T327 | 0.0895 | 0.0414 | 0.0296 | 0.0185 | 0.0006 |
| M436T365 | 2.4024 | 0.3154 | 0.1681 | 0.0076 | 0.0051 |

**Table S4. At two postoperative time points, the contents of 92 plasma metabolites all showed significant changes**

| **Metabolite** | **MEAN**  **(pre)** | **MEAN**  **(post 6 month)** | **MEAN**  **(post 12 month)** | **P-VALUE**  **(pre vs post 6 month)** | **P-VALUE**  **(pre vs post 12 month)** |
| --- | --- | --- | --- | --- | --- |
| 3-methylcytidine | 0.0016 | 0.0031 | 0.0029 | 0.0500 | 0.0272 |
| 6-Hydroxynorleucine | 0.0254 | 0.0333 | 0.0372 | 0.0484 | 0.0028 |
| Hyodeoxycholic acid | 0.1166 | 0.0653 | 0.0413 | 0.0429 | 0.0045 |
| Carnitine | 7.8293 | 9.1940 | 9.7274 | 0.0447 | 0.0024 |
| Topiramate | 7.5255 | 24.8504 | 20.1791 | 0.0013 | 0.0265 |
| SM(d18:1/18:0) | 0.3106 | 0.3960 | 0.5512 | 0.0216 | 0.0000 |
| 10,13-dimethyl-17-oxo-3-sulfooxy-1,2,3,4,7,8,9,11,12,14,15,16-dodecahydrocyclopenta[a]phenanthrene | 1.7953 | 13.5877 | 12.2256 | 0.0059 | 0.0425 |
| Ursodeoxycholic acid | 0.1166 | 0.0653 | 0.0413 | 0.0429 | 0.0045 |
| 5-Methylcytidine | 0.0016 | 0.0031 | 0.0029 | 0.0500 | 0.0272 |
| Adipic acid | 0.0392 | 0.0295 | 0.0246 | 0.0270 | 0.0014 |
| 2-Methylglutaric acid | 0.0392 | 0.0295 | 0.0246 | 0.0270 | 0.0014 |
| Rhamnose | 0.0283 | 0.0368 | 0.0414 | 0.0133 | 0.0001 |
| SM(d18:1/17:0) | 0.0417 | 0.0538 | 0.0736 | 0.0194 | 0.0004 |
| GlcCer(d18:1/16:0) | 0.0135 | 0.0182 | 0.0208 | 0.0131 | 0.0065 |
| 3-Methylglutaric acid | 0.0392 | 0.0295 | 0.0246 | 0.0270 | 0.0014 |
| 3-Hydroxyisovaleric acid | 0.6446 | 0.4025 | 0.4297 | 0.0080 | 0.0065 |
| LPI(18:2) | 0.0287 | 0.0163 | 0.0163 | 0.0037 | 0.0043 |
| Valnoctamide | 0.0119 | 0.0103 | 0.0098 | 0.0323 | 0.0109 |
| Valpromide | 0.0119 | 0.0103 | 0.0098 | 0.0323 | 0.0109 |
| Nomilin | 0.0106 | 0.0046 | 0.0056 | 0.0269 | 0.0415 |
| 5.alpha.-Androstan-3.beta.-ol-17-one glucosiduronate | 0.0007 | 0.0048 | 0.0046 | 0.0317 | 0.0216 |
| 1,4-Bis(4-morpholinylmethyl)-2,3-naphthalenediol | 0.0329 | 0.0139 | 0.0129 | 0.0135 | 0.0104 |
| N-Nervonoyl-D-erythro-sphingosylphosphorylcholine | 0.8074 | 1.0594 | 1.1088 | 0.0245 | 0.0070 |
| 4-Methyl-1H-pyrazole | 0.0230 | 0.0326 | 0.0263 | 0.0033 | 0.0011 |
| PC 8:0_22:3 | 0.0038 | 0.0025 | 0.0016 | 0.0314 | 0.0003 |
| D-myo-Inositol, 1-[2-hydroxy-3-[(1-oxo-9,12-octadecadienyl)oxy]propyl hydrogen phosphate], [S-(Z,Z)]- | 1.1534 | 0.8235 | 0.6708 | 0.0198 | 0.0017 |
| SMGDG O-8:0_16:4 | 0.0149 | 0.0061 | 0.0069 | 0.0014 | 0.0043 |
| MGDG O-20:5_3:0 | 0.0898 | 0.0474 | 0.0270 | 0.0058 | 0.0000 |
| 2-(1H-Pyrrol-1-yl)benzoic acid | 0.0144 | 0.0633 | 0.0695 | 0.0027 | 0.0347 |
| Lyso-sphingomyelin | 0.0032 | 0.0045 | 0.0063 | 0.0167 | 0.0001 |
| n-Octadecylamine | 0.0038 | 0.0043 | 0.0056 | 0.0384 | 0.0010 |
| N-(1.3-dihydroxyoctadeca-4.14-dien-2-yl)palmitamide | 0.0011 | 0.0027 | 0.0035 | 0.0027 | 0.0167 |
| 3-[5-(4-Pyridinyl)-4H-1,2,4-triazol-3-yl]pyridine | 0.0037 | 0.0076 | 0.0077 | 0.0038 | 0.0000 |
| 2-Amino-6-(methylsulfanyl)-3,5-pyridinedicarbonitrile | 0.1891 | 0.2335 | 0.2255 | 0.0088 | 0.0205 |
| 2-Chloro-5-(4H-1,2,4-triazol-4-yl)benzoic acid | 0.0098 | 0.0368 | 0.0282 | 0.0010 | 0.0227 |
| Dodeca-2(E),4(E)-dienoic acid | 0.0711 | 0.1841 | 0.2269 | 0.0044 | 0.0000 |
| Trihexosylceramide_(d18:1/16:0) | 0.0002 | 0.0003 | 0.0003 | 0.0269 | 0.0001 |
| N-[(4E,8E)-1,3-dihydroxyoctadeca-4,8-dien-2-yl]hexadecanamide | 0.0191 | 0.0313 | 0.0370 | 0.0001 | 0.0022 |
| 18-Carboxydinorleukotriene B4 | 0.2062 | 0.0927 | 0.1041 | 0.0014 | 0.0048 |
| 4-Amino-3-bromobenzenesulfonamide | 0.1065 | 0.1132 | 0.1130 | 0.0481 | 0.0440 |
| Alpinumisoflavone | 0.0202 | 0.0248 | 0.0231 | 0.0008 | 0.0084 |
| (3'-sulfo)Gal.beta.-Cer(d18:1/22:0) | 0.0051 | 0.0068 | 0.0074 | 0.0352 | 0.0007 |
| SM 24:0 | 0.0073 | 0.0140 | 0.0116 | 0.0029 | 0.0101 |
| 6-(Trifluoromethyl)-1H-pyrazolo[3,4-d]pyrimidin-4-ol | 0.1794 | 0.2291 | 0.2175 | 0.0006 | 0.0139 |
| Phe-Met-Arg-Phe-amide | 0.0564 | 0.0379 | 0.0298 | 0.0195 | 0.0022 |
| (2R)-2-(2,5-Difluorophenyl)pyrrolidine | 0.2112 | 0.2866 | 0.3037 | 0.0340 | 0.0031 |
| 4-tert-Butyl-N'-[(E)-(2-hydroxynaphthalen-1-yl)methylidene]benzohydrazide | 0.2057 | 0.0737 | 0.0524 | 0.0017 | 0.0005 |
| 3-Cyclohexyl-2-sulfanyl-4(3H)-quinazolinone | 0.0086 | 0.0152 | 0.0190 | 0.0195 | 0.0229 |
| 1-Hydroxy-2-naphthoic acid | 2.6606 | 2.0021 | 1.9147 | 0.0138 | 0.0335 |
| Glyceryl trimyristate | 0.0125 | 0.0163 | 0.0305 | 0.0264 | 0.0002 |
| 2-Chloro-5,6,7,8-tetrahydroquinoxaline | 0.0731 | 0.0957 | 0.0929 | 0.0013 | 0.0071 |
| LPG O-17:1 | 0.0773 | 0.0366 | 0.0228 | 0.0037 | 0.0003 |
| PI 4:0_20:4 | 0.0095 | 0.0028 | 0.0043 | 0.0065 | 0.0288 |
| 4-Phenyl-5-(2-phenylethyl)-4H-1,2,4-triazol-3-yl hydrosulfide | 0.0354 | 0.1386 | 0.1222 | 0.0167 | 0.0033 |
| 6-Amino-2,4-dichloro-3-ethylphenol | 0.0026 | 0.0093 | 0.0071 | 0.0015 | 0.0236 |
| N-(5-Chloro-2-methoxyphenyl)-N'-(5-methyl-3-isoxazolyl)urea | 0.0313 | 0.1138 | 0.0866 | 0.0010 | 0.0241 |
| N,N-Dimethylformamide | 0.0396 | 0.0557 | 0.0613 | 0.0087 | 0.0031 |
| LNAPS 2:0/N-16:3 | 0.0073 | 0.0038 | 0.0019 | 0.0364 | 0.0027 |
| (1R,4R,6R)-4-Amino-2-oxabicyclo[3.1.0]hexane-4,6-dicarboxylic acid | 0.6089 | 0.4643 | 0.4369 | 0.0161 | 0.0264 |
| (25S)-7-Dafachronic acid | 0.0046 | 0.0025 | 0.0015 | 0.0113 | 0.0002 |
| Unguisin E | 0.0057 | 0.0019 | 0.0023 | 0.0024 | 0.0056 |
| PC(16:0/P-16:0) | 0.0852 | 0.1178 | 0.1075 | 0.0378 | 0.0067 |
| PC(14:0/P-18:1(9Z)) | 0.0181 | 0.0279 | 0.0247 | 0.0242 | 0.0028 |
| 5-Sulfosalicylic acid | 0.0955 | 0.0438 | 0.0321 | 0.0480 | 0.0216 |
| LNAPS 3:0/N-13:0 | 0.2768 | 0.1877 | 0.1330 | 0.0173 | 0.0001 |
| LNAPE 3:0/N-24:7 | 0.0952 | 0.0201 | 0.0229 | 0.0233 | 0.0276 |
| Dictamnine | 0.0801 | 0.1171 | 0.1237 | 0.0113 | 0.0038 |
| SM(d16:1/24:1(15Z)) | 0.6326 | 0.8414 | 0.9792 | 0.0040 | 0.0031 |
| DGDG O-10:0_3:0 | 0.1488 | 0.0464 | 0.0334 | 0.0027 | 0.0002 |
| SM(d17:1/24:1(15Z)) | 0.1507 | 0.2022 | 0.2694 | 0.0042 | 0.0000 |
| Stigmastane-3,6-dione | 0.5865 | 0.8033 | 0.8253 | 0.0126 | 0.0035 |
| 1,3-Benzothiazol-2-yl difluoromethyl sulfide | 0.0018 | 0.0056 | 0.0045 | 0.0017 | 0.0320 |
| Palitantin | 0.0029 | 0.0023 | 0.0023 | 0.0482 | 0.0331 |
| 1,2,4-Oxadiazole, 5-[4-phenyl-5-(trifluoromethyl)-2-thienyl]-3-[3-(trifluoromethyl)phenyl]- | 0.0061 | 0.0209 | 0.0163 | 0.0011 | 0.0286 |
| Benzenemethanaminium, N-cyclopentyl-N,N-dimethyl-3,4-bis(tetradecyloxy)- | 0.0062 | 0.0120 | 0.0152 | 0.0006 | 0.0002 |
| taxifolin | 0.0208 | 0.0771 | 0.0591 | 0.0009 | 0.0207 |
| (2-{[2-(2,6-Dimethylphenoxy)ethyl]sulfanyl}-1H-benzimidazol-1-yl)acetic acid | 0.2480 | 0.9154 | 0.7022 | 0.0011 | 0.0240 |
| PA 3:0_14:1 | 0.0434 | 0.0290 | 0.0215 | 0.0134 | 0.0002 |
| Isoginkgetin | 0.0026 | 0.0030 | 0.0028 | 0.0055 | 0.0245 |
| 2-Phenyl-4-(1-pyrrolidinyl)quinazoline | 0.1162 | 0.1261 | 0.1252 | 0.0391 | 0.0276 |
| Ethyl 2-(4-(t-BOC)piperazin-1-yl)-4-hydroxypyrimidine-5-carboxylate | 0.0062 | 0.0300 | 0.0262 | 0.0013 | 0.0309 |
| 2''-O-Methylmitorubrin | 0.0444 | 0.0520 | 0.0489 | 0.0002 | 0.0496 |
| 5'-Methoxybilobetin | 0.0010 | 0.0012 | 0.0011 | 0.0079 | 0.0340 |
| 15-Ketofluprostenol isopropyl ester | 0.0298 | 0.0172 | 0.0163 | 0.0493 | 0.0424 |
| Alachlor | 0.0006 | 0.0003 | 0.0003 | 0.0132 | 0.0048 |
| M495T460 | 0.0290 | 0.0157 | 0.0075 | 0.0325 | 0.0016 |
| M611T408 | 0.0141 | 0.0046 | 0.0029 | 0.0055 | 0.0018 |
| M482T327 | 0.0003 | 0.0021 | 0.0018 | 0.0271 | 0.0001 |
| M505T400 | 0.0023 | 0.0033 | 0.0036 | 0.0421 | 0.0183 |
| M480T383 | 0.0092 | 0.0103 | 0.0100 | 0.0228 | 0.0253 |
| M649T412 | 0.0036 | 0.0022 | 0.0009 | 0.0306 | 0.0000 |
| M264T310 | 0.0489 | 0.1808 | 0.1376 | 0.0010 | 0.0244 |

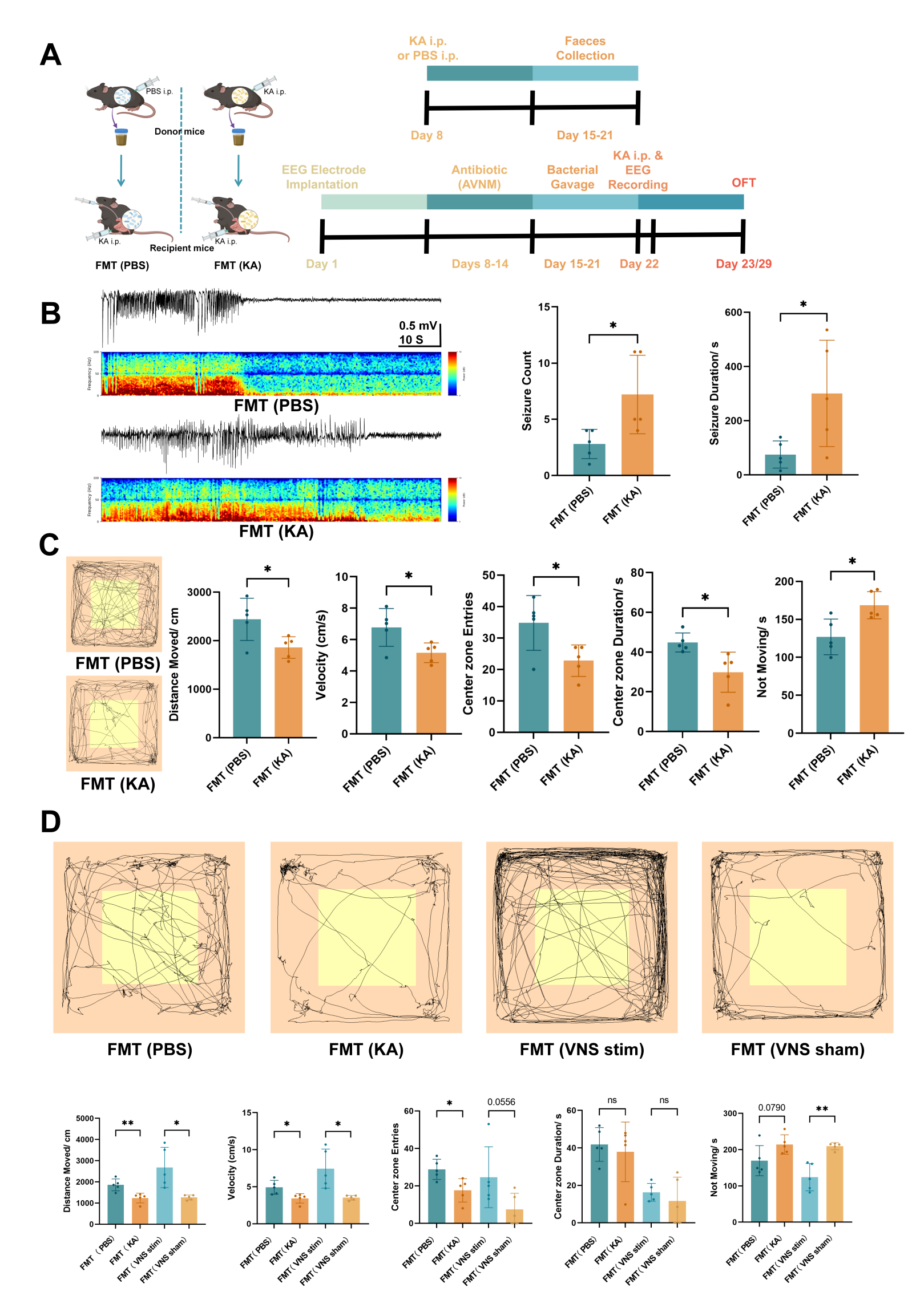

**Figure S2. Epilepsy-Associated Microbiota Exacerbates Seizure Susceptibility in Recipient Mice, Related to Figure 2**

(A) Experimental schematic. Antibiotic-treated recipient mice received FMT from donors injected with PBS (FMT-PBS, n=5) or KA (FMT-KA, n=5) prior to seizure induction.
(B) Representative EEG traces (left) and quantification (right). The FMT-KA group exhibited a higher frequency and longer duration of seizures compared to the FMT-PBS group.
(C) Representative track plots from the OFT 1 day post-induction. The FMT-PBS group showed greater locomotor activity (distance, velocity), center zone entries, and center zone duration, with reduced immobility time compared to the FMT-KA group.
(D) Representative track plots from the OFT 1 week post-induction. Consistent with day 1, the FMT-PBS group maintained higher activity and exploration levels with lower immobility compared to the FMT-KA group. Furthermore, in the VNS cohort (described in Figure 2), the FMT-VNS group maintained higher locomotor activity and center zone entries with reduced immobility compared to the FMT-Sham group at this time point.

Data are represented as mean ± SEM with individual data points shown (B, C, and D). Statistical significance was determined by unpaired t-test. *p < 0.05, **p < 0.01, ***p < 0.001, ****p < 0.0001; ns, not significant. **FMT**, Fecal microbiota transplantation; **PBS**, Phosphate-buffered saline; **KA**, Kainic acid; **i.p.**, Intraperitoneal; **EEG**, Electroencephalogram; **AVNM**, Antibiotic cocktail; **OFT**, Open field test; **VNS**, Vagus nerve stimulation.

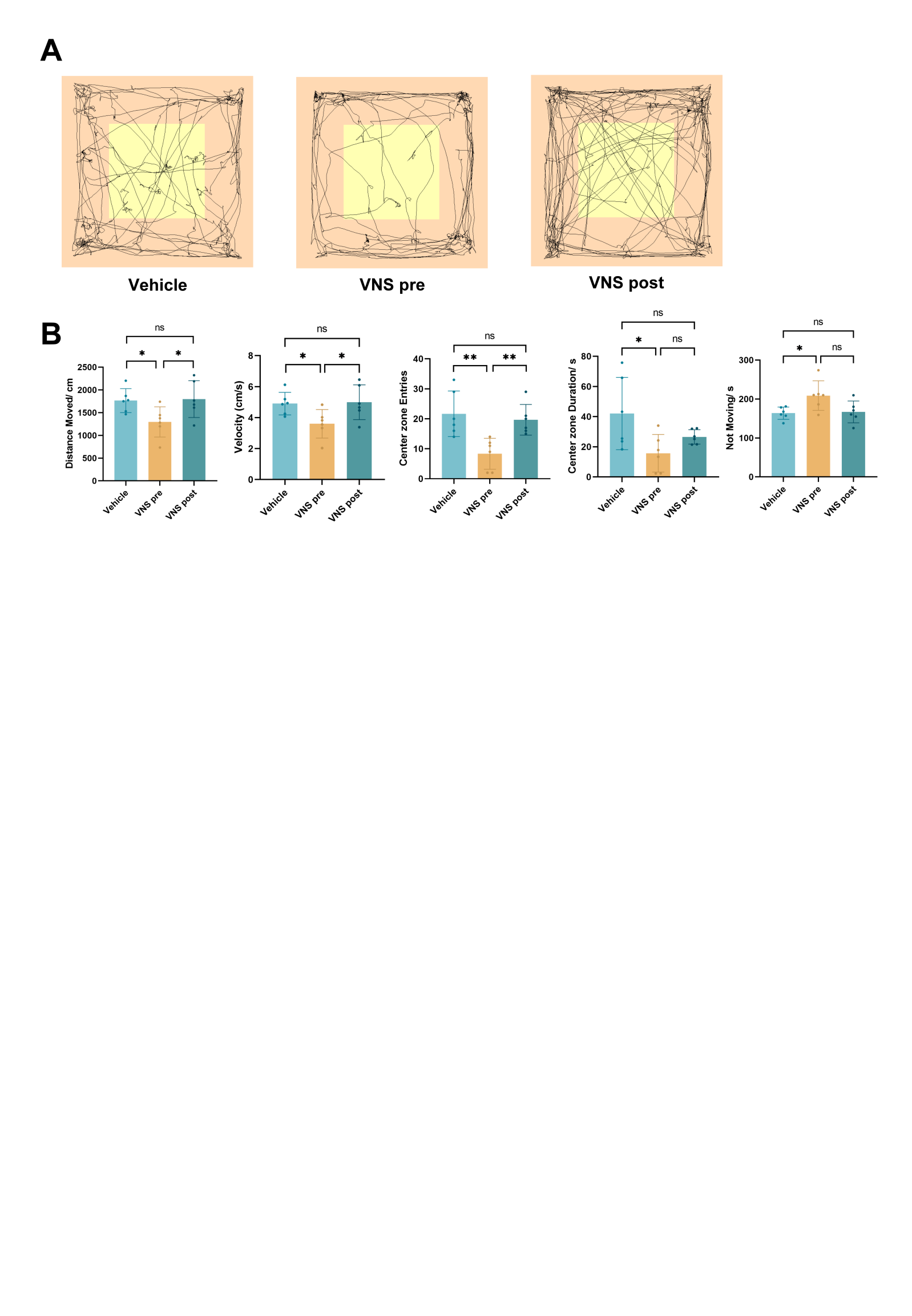

**Figure S3. The protective effect of the gut microbiota from VNS patients on recipient mice persisted for one week after modeling.**

**(A, B)** Open-field test conducted one week after epilepsy induction. In the VNS post group (n = 6), distance moved, velocity, and center-zone entries were higher than in the VNS pre group (n = 6).

Data are presented as mean ± standard error, and individual data points are shown (B). Unpaired t-test (B). *p < 0.05, **p < 0.01, ***p < 0.001, ****p < 0.0001, ns, no significant difference. **VNS**, Vagus nerve stimulation.

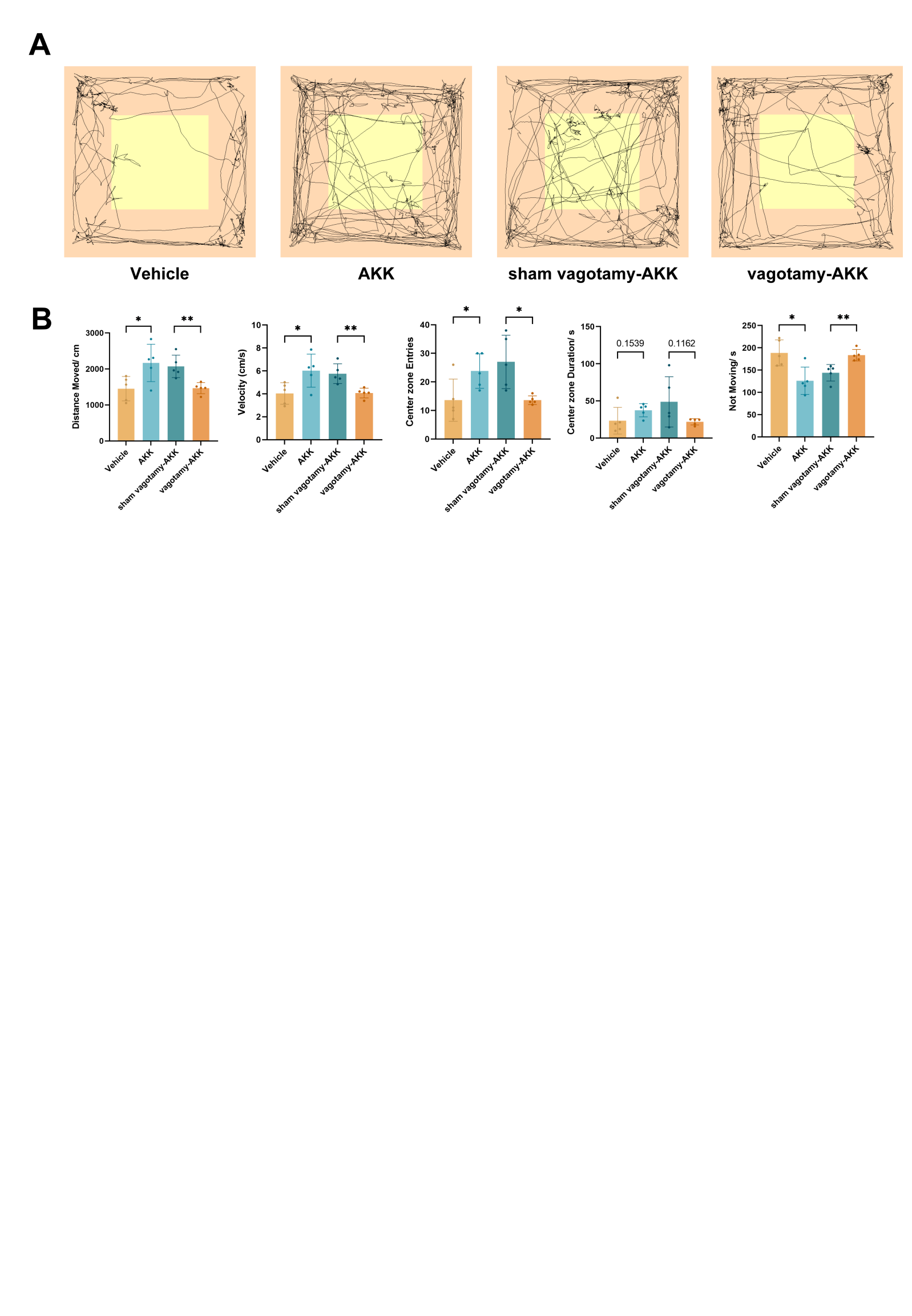
**Figure S4. The protective effect of Akkermansia muciniphila on recipient mice persisted for one week after modeling.**

**(A, B)** Open-field test one week after epilepsy induction. The AKK group (n = 6) exhibited greater distance moved, higher mean velocity, and more center-zone entries, with reduced immobility time, compared with the Vehicle group (n = 6). The vagotomy-AKK group (n = 6) showed lower distance moved, slower velocity, fewer center-zone entries, and longer immobility time than the sham vagotomy-AKK group (n = 6).

Data are presented as mean ± standard error, and individual data points are shown (B). Unpaired t-test (B). *p < 0.05, **p < 0.01, ***p < 0.001, ****p < 0.0001, ns, no significant difference. **AKK**, *Akkermansia muciniphila*.

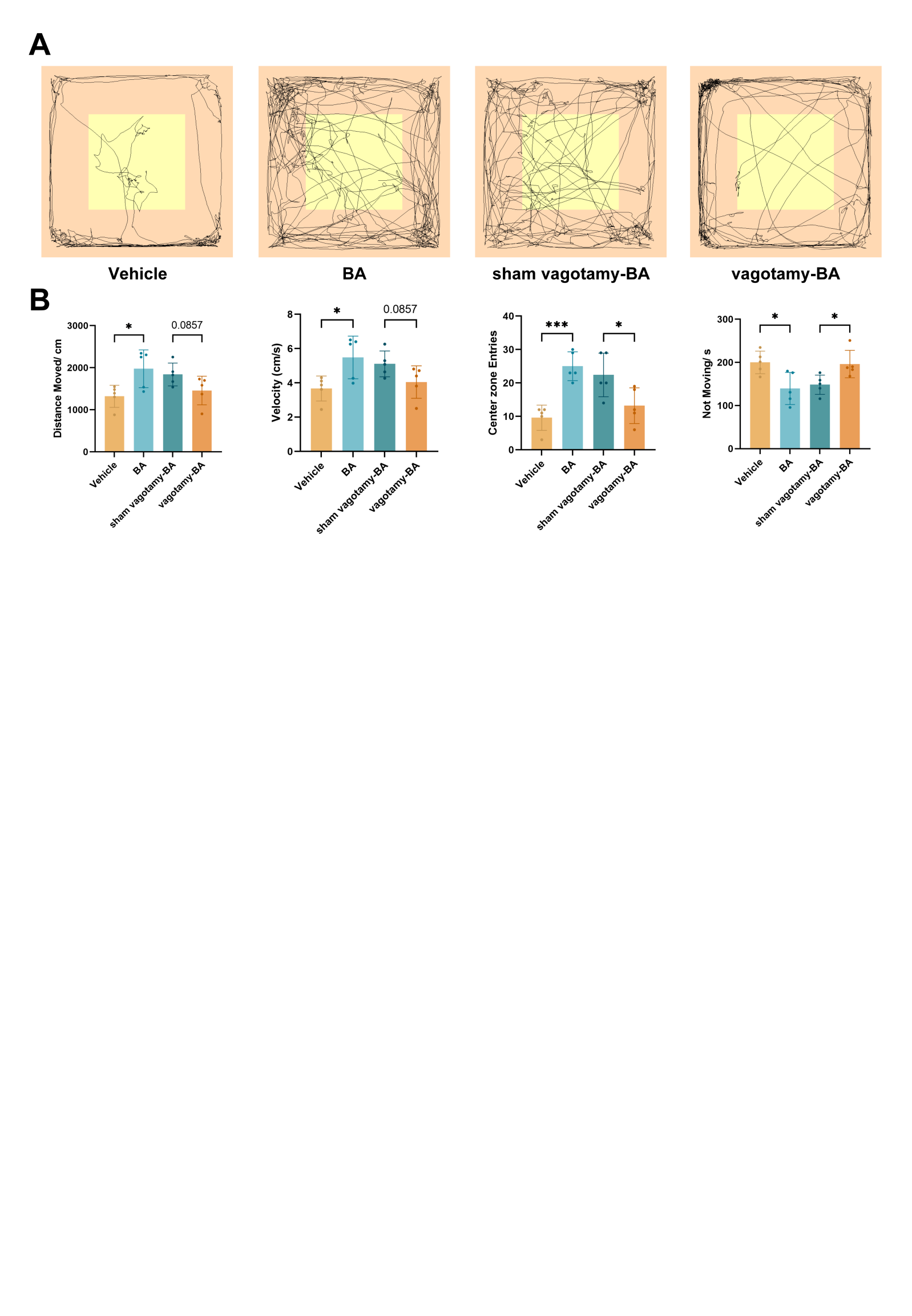

**Figure S5. The protective effect of butyrate on recipient mice persisted for one week after modeling.**

**(A, B)** Open-field test one week after epilepsy modeling. The BA group (n = 5) exhibited greater distance moved, higher velocity, more center-zone entries, and reduced immobility time compared with the Vehicle group (n = 5). The vagotomy-BA group (n = 5) showed lower distance moved, slower velocity, fewer center-zone entries, and longer immobility time than the sham vagotomy-BA group (n = 5).

Data are presented as mean ± standard error, and individual data points are shown (B). Unpaired t-test (B). *p < 0.05, **p < 0.01, ***p < 0.001, ****p < 0.0001, ns, no significant difference. **BA**, butyrate.
